# Analysis of 14q12 microdeletions reveals novel regulatory loci for the neurodevelopmental disorder-related gene, *FOXG1*

**DOI:** 10.1101/2025.04.11.648472

**Authors:** Aishwarya Ramamurthy, Masha D Bandouil, Likhita Aluru, Ojasi Joshi, Esther Yoon, Nicholas Bodkin, Jennifer Z Cheng, Carina G Biar, Jeffrey D Calhoun, Gemma L Carvill

## Abstract

Up to 17% of neurodevelopmental disorders (NDDs) can be explained by pathogenic structural variants (SVs) that disrupt coding regions and elicit gene dosage defects. However, noncoding SVs which can perturb *cis*-regulatory elements (CREs) and downstream gene expression are understudied. In this study, we describe multiple 14q12 deletions downstream of NDD-related gene *FOXG1* in individuals with overlapping phenotypes of *FOXG1* haploinsufficiency. We show that deletion of a minimum region of overlap (MRO) reduced *FOXG1* expression, disrupted CREs and altered *FOXG1*’s native genomic interactions. Deleting the MRO did not fully eliminate *FOXG1* expression, indicating that multiple CREs likely cooperate to regulate *FOXG1* and would need to be deleted to completely prevent expression. The transcriptomic profiles of MRO loss overlap in part with *FOXG1* loss, including direct FOXG1 targets, indicating converging molecular pathways. These findings expand the scope of *FOXG1’s* complex regulatory region, and more broadly, of regulatory SVs in NDD susceptibility.

## Introduction

Neurodevelopmental disorders (NDDs) are a heterogenous collection of developmental disorders that impair functions of the brain and the nervous system. These disorders have childhood onset and include, among others, intellectual disability (ID), autism spectrum disorder (ASD) and epilepsy. It is now well established that disease-causing protein-coding variants in hundreds of genes account for up to 40% of NDDs ^1,2^. This still leaves a large fraction of cases unexplained, rekindling interest in investigating the noncoding regulatory regions of the genome for causative variants in individuals with NDDs.

In particular, structural variants (SVs) are attractive candidates for understanding noncoding variation due to the thousands to millions of base pairs that can be disrupted and thus, in theory, easier interpretation as compared to single nucleotide variants (SNVs). SVs, defined as alterations greater than 50 base pairs, include deletions, duplications, insertions, inversions, and translocations. SVs can overlap known NDD-related genes and cause these conditions through gene dosage effects. Coding SVs account for 15-17% of NDDs in general ^3,4^ and 10-12% of specific NDDs, such as the genetic epilepsies ^5–7^. However, some SVs do not span coding regions but rather disrupt cis-regulatory elements (CREs) or higher order chromatin structures, including topologically associated domains (TADs). When SVs disrupt CREs, promoter-enhancer interactions can be perturbed, which affects gene transcription ^8–10^. For example, noncoding SNVs and duplications in the *GATA2* regulatory region cause hereditary congenital facial paresis ^11^. These variants cause disease by aberrant prolonged *GATA2* expression which in turn impairs an essential neural fate switch during development ^11^. SVs can also disrupt TAD boundaries, leading to an alteration of the native chromatin architecture, promoting non-canonical promoter-enhancer interactions and gene misexpression. TAD disruption by SVs has been reported in human limb malformations and congenital disorders ^12,13^. Moreover, noncoding variants in the TAD encompassing NDD-related gene, *MEF2C* were present in excess in a large genomic screen of individuals with NDDs, and deletion of a proximal *MEF2C* TAD boundary resulted in reduced gene expression ^14,15^. Collectively, these studies highlight an emerging role for noncoding SVs in the etiology of NDDs. Here we add to this burgeoning field by describing NDD-related microdeletions at the 14q12 locus flanking the gene, *FOXG1*.

*FOXG1* encodes Forkhead Box G1, a key transcription factor in forebrain development that is highly dosage sensitive ^16^. *FOXG1* expression is especially critical during corticogenesis, triencephalon specification and development, as well as maintenance of mature neurons ^17,18^. Neurodevelopment is reliant not only on precise spatio-temporal *FOXG1* expression, but also on its cellular expression levels; both increased and decreased *FOXG1* expression is associated with poor neurological outcomes ^16,19,20^. Heterozygous loss of *FOXG1* causes *FOXG1* syndrome, a congenital NDD characterized by epilepsy, ID, microcephaly, developmental delays, loss of motor skills and overlapping clinical features with Rett syndrome ^21–23^. Several heterozygous *de novo* whole gene deletions, truncating variants and loss of function missense variants cause *FOXG1* syndrome. The pathogenic mechanism is thus haploinsufficiency, wherein one autosomal allele of *FOXG1* is functional, but insufficient to prevent disease ^22^. In addition, *de novo* duplications of *FOXG1* result in increased gene dosage and are associated with developmental epilepsy, ID, and severe speech impairment ^24^.

SVs overlapping putative CREs have also been described in individuals with overlapping clinical features of *FOXG1* haploinsufficiency, but without coding variants ^23,25–27^. While the impact of these deletions on *FOXG1* expression was examined, results have been contradictory, likely owing to the virtually absent *FOXG1* expression in accessible peripheral patient biospecimens (Supplementary Fig. 1). Specifically, one group showed increased *FOXG1* in blood and saliva from individuals with 14q12 deletions and predicted a single region of overlap (SRO) to be regulatory ^25^. Conversely, another showed reduced *FOXG1* expression in blood, despite shared regions of overlap across patient-specific deletions (Fig. 1) ^26^. To address these inconsistencies, here we refined the SRO to a new minimum region of overlap (MRO) in individuals with *FOXG1* haploinsufficient clinical features and downstream deletions, and modeled the MRO in the HAP1 cell line which robustly expresses *FOXG1*. We refine the regulatory landscape of the *FOXG1* locus and demonstrate that MRO deletion reduced *FOXG1* expression, likely contributing to haploinsufficiency in individuals with NDDs.

**Figure 1:**
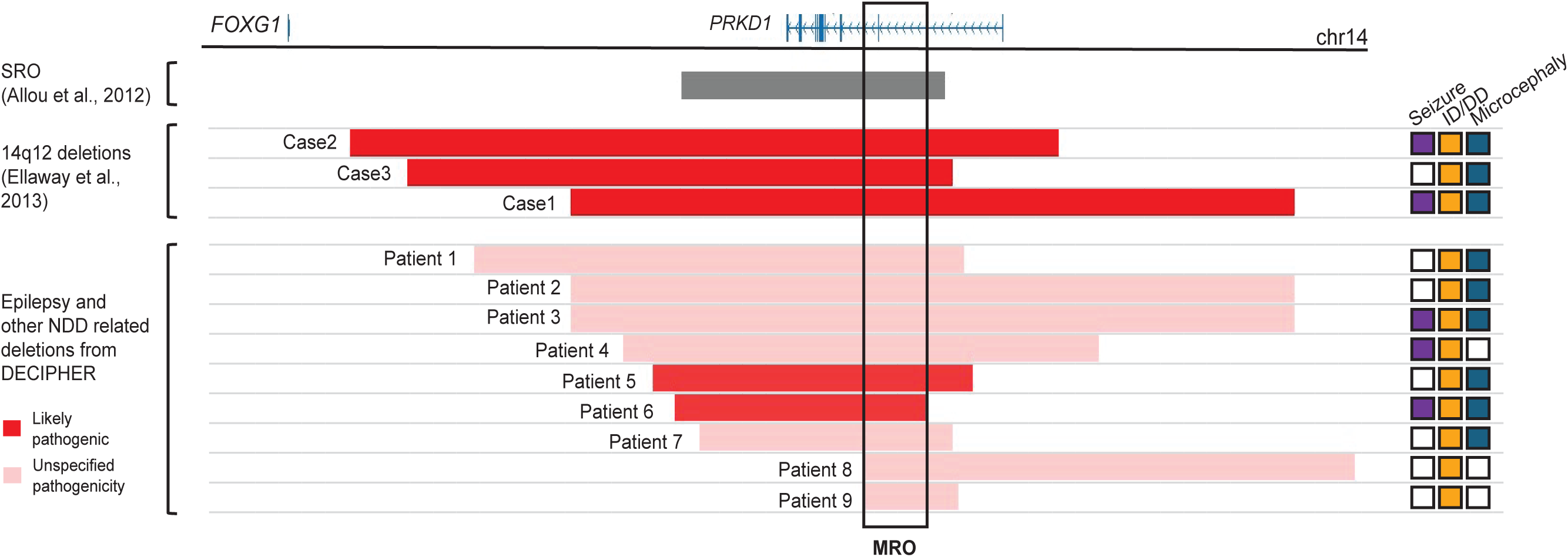
14q12 microdeletions downstream of *FOXG1* in this study. Deletions present in individuals with epilepsy and other NDDs in DECIPHER are represented by horizontal bars and color-coded for pathogenicity. Additionally, three deletions from prior work that met our selection criteria are also shown (red horizontal bars) ^26^. The overlap between the 14q12 deletions was used to define the minimum region of overlap (MRO), in this case by the deletion breakpoints in patients 6, 8 and 9 (black enclosure). The panel on the right indicates whether individuals with the respective 14q12 deletions presented with seizures, intellectual disability (ID) or developmental delays (DD) and microcephaly. Grey bar represents a single overlapping region (SRO) in the 14q12 region previously predicted to regulate *FOXG1* ^25^.

## Results

### Deletion of downstream *FOXG1* 14q12 minimum region of overlap leads to reduced *FOXG1* expression

We identified 9 microdeletions in the 14q12 region downstream of *FOXG1* described in DECIPHER. These deletions were reported in individuals (6 females and 3 males) with ID (n=8), seizures (n=3) and microcephaly (n=6) (Fig. 1 and Table 1). Most of these deletions were *de novo* (patients 2-7); one deletion was maternally inherited (patient 9) and two were of unknown inheritance (patient 1 and 8). None of these deletions encompassed any other gene except *PRKD1. PRKD1* is tolerant to loss of function in the general population with a pLI score of 0 and LOEUF of 0.76 (gnomAD, ^28,29^), has very low expression in the brain compared to other tissues (GTEx), and the mouse knockout has no neurological phenotype (MGI:99879), collectively suggesting *PRKD1* loss is not associated with NDDs. Thus, we hypothesized that disruption of CREs in the region likely contributed to the phenotype via disruption of *FOXG1* expression. We selected the minimum region of overlap (MRO) between these deletions for modeling in the HAP1 cell line. This region (hg38: chr14:29,704,736-29,803,897) spans ∼0.1 Mb and is ∼0.93 Mb downstream of *FOXG1*. The MRO was examined for variants in the general population using gnomAD and no large deletions were observed in this locus (Supplementary Fig. 2).

**Table 1:** Genetic and clinical details of 14q12 heterozygous deletion variants in individuals with epilepsy, ID and/or microcephaly considered in defining the minimum region of overlap (MRO)

| Patient | Decipher ID | Sex | Deletion coordinates (hg38) | CREs deleted (Fig. 3) | Inheritance | Pathogenicity assignment in DECIPHER | Seizures | Intellectual Disability (ID) | Microcephaly | Additional clinical features |
| --- | --- | --- | --- | --- | --- | --- | --- | --- | --- | --- |
| DECIPHER |  |  |  |  |  |  |  |  |  |  |
| 1 | 260836 | 46XY | 14:29068799-29865858 | 6 VISTA enhancers, EC, HAP1-CRE1 | Unknown | Unspecified | - | Yes | Yes |  |
| 2 | 248405 | 46XY | 14:29226052-30403168 | 7 VISTA enhancers, EC, HAP1-CRE1 | *De novo* | Unspecified | - | Yes | Yes |  |
| 3 | 3810 | 46XX | 14:29226052-30403168 | 7 VISTA enhancers, EC, HAP1-CRE1 | *De novo* | Unspecified | Yes | Yes | Yes | Bruxism, hypoplasia of the corpus callosum, delayed speech and language development |
| 4 | 258584 | 46XX | 14:29312198-30083730 | 3 VISTA enhancers, EC, HAP1-CRE1 | *De novo* | Unspecified | Yes | Yes | - | Atypical behavior, delayed speech and language development |
| 5 | 358785 | 46XX | 14:29360119-29879052 | 3 VISTA enhancers, EC, HAP1-CRE1 | *De novo* | Likely Pathogenic | - | Yes | Yes (secondary) | Agenesis of corpus callosum |
| 6 | 414554 | Female | 14:29395780-29803897 | 1 VISTA enhancer, EC, HAP1-CRE1 | *De novo* | Likely Pathogenic | Yes | - | Yes | Delayed myelination, global developmental delay, simplified gyral pattern |
| 7 | 252353 | 46XX | 14:29435514-29847454 | 1 VISTA enhancer, EC, HAP1-CRE1 | *De novo* | Unspecified | - | Yes | Yes | Hypoplasia of the corpus callosum |
| 8 | 259935 | 46XX | 14:29704706-30500956 | 2 VISTA enhancers, HAP1-CRE1 | Unknown | Unspecified | - | Yes | - | Atypical behavior |
| 9<sup>27</sup> | 284715 | 46XY | 14:29704736-29856618 | HAP1-CRE1 | Maternally inherited | Unspecified | - | Yes (mild) | - | Delayed speech and language development |
| Published (Ellaway *et al.*)<sup>26</sup> |  |  |  |  |  |  |  |  |  |  |
| Case 1 | - | Female | 14:29226052-30403168 | 7 VISTA enhancers, EC, HAP1-CRE1 | *De novo* | - | Yes | - | Yes | Intellectual impairment, agenesis of corpus callosum, global developmental delay |
| Case 2 | - | Female | 14:28868223-30018792 | 6 VISTA enhancers, EC, HAP1-CRE1 | *De novo* | - | Yes | - | Yes | Intellectual impairment, agenesis of corpus callosum, global developmental delay |
| Case 3 | - | Male | 14:28960335-29847513 | 6 VISTA enhancers, EC, HAP1-CRE1 | *De novo* | - | - | - | Yes | Intellectual impairment, severe developmental delay |

Abbreviations: EC - Enhancer Cluster

We selected the HAP1 cell line due to its haploid nature, which facilitates genome editing, as well as the endogenous expression of *FOXG1* (Supplementary Fig. 1 and Fig. 2). We successfully deleted the MRO using dual flanking guides in HAP1s (14q12-MRO-Del); the entire *FOXG1* coding region was similarly deleted as a positive control (FOXG1^KO^). We verified the genotypes of 14q12-MRO-Del (deletion of hg38: chr14:29,704,715-29,803,900) and FOXG1^KO^ (deletion of hg38: chr14:28,767,269-28,768,758) using Sanger sequencing, and in the 14q12-MRO-Del confirmed the absence of major off-target SVs using genome sequencing (Supplementary Fig. 3). We confirmed the loss of *PRKD1* exon 2 and flanking intronic sequence at both the genomic DNA and mRNA level in 14q12-MRO-Del (Supplementary Fig. 3-4).

**Figure 2:**
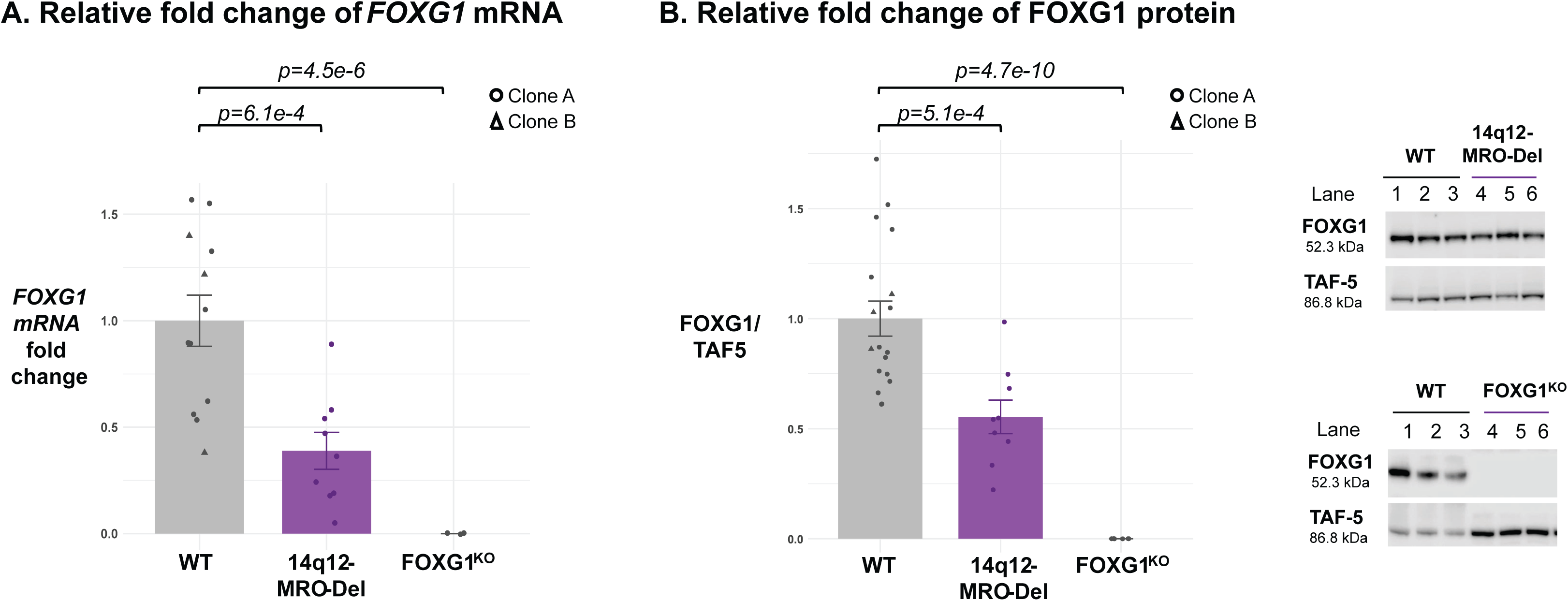
*FOXG1* expression is reduced with the loss of the MRO locus (A) *FOXG1* mRNA fold change relative to *EAR* assayed by qRT-PCR (n: WT Clone A = 9, WT Clone B = 3, 14q12-MRO-Del = 9, FOXG1^KO^ = 3) (Two sample T-test, WT vs 14q12-MRO-Del, p= 6.1e-4; WT vs FOXG1^KO^, p = 4.5e-6). (B) FOXG1 protein levels relative to TAF5 were assayed using Western Blotting (n: WT Clone A =15, WT Clone B = 3, 14q12-MRO-Del = 9, FOXG1^KO^ = 6) (Two sample T-test, WT vs 14q12-MRO-Del, p= 5.1e-4; WT vs FOXG1^KO^, p = 4.7e-10). Error bars represent one standard error.

Using qRT-PCR, we demonstrated complete loss of *FOXG1* mRNA expression in FOXG1^KO^ and a 61% reduction in the 14q12-MRO-Del line (p-value 0.0006, Welch’s T test) (Fig. 2A). At the protein level, western blotting revealed similar loss of protein in FOXG1^KO^ and a 45% reduction in the 14q12-MRO-Del (p-value 0.0005, Welch’s T test) (Fig. 2B). Collectively, these results demonstrate that deleting the MRO reduces *FOXG1* expression, though not to the same degree as deletion of the complete coding region.

### Deletion of the MRO locus alters *FOXG1’s* native contacts with downstream CREs

To examine the causal mechanism of reduced *FOXG1* expression with MRO deletion, we examined chromatin interactions of the *FOXG1* locus with downstream CREs in wildtype and 14q12-MRO-Del lines using Unique Molecular Identifier (UMI)-4C. We anchored the viewpoint proximal to *FOXG1’s* promoter (Supplementary Table 4) and identified 9997 UMIs in WT and 6805 UMIs in 14q12-MRO-Del. UMI-4C contact intensity profiles were generated which highlight multiple promoter – CRE interactions across the 14q12 region (Fig. 3). The overwhelming majority of contacts lie downstream of *FOXG1*, with very few contacts, and no peaks upstream. The CRE landscape across this region is complex with many known and predicted regulatory regions and most have high levels of conservation (Fig. 3). This includes nine VISTA enhancers active in neurodevelopment, of which six show forebrain-specific activity (Hs1064, Hs566, Hs1523, Hs342, Hs433, Hs344) (Supplementary Fig. 5) ^30,31^. In addition, there is an enhancer cluster near Hs598, with five active forebrain-specific enhancers in a zebrafish model ^32^. A massively parallel reporter assay (MPRA) in fetal brain and cerebral organoids further reveals dozens of active CREs ^33^. As well as these demonstrated active CREs, in both HAP1s and neural cell lineages there are a number of putative enhancers, defined by the histone modification, H3K27Ac ^30,31,34–37^. Deleting the MRO not only led to loss of UMI-4C contacts in this region as expected, but also altered native *FOXG1* genomic interactions. Specifically, significant differences in UMI contacts between WT and 14q12-MRO-Del were observed at six 14q12 loci, including four regions with increased contacts and two that were decreased (Fig. 3, Supplementary Fig. 6). Of note, four of these enhanced contacts (3 increased, one decreased) were located 1.5 Mb downstream of *FOXG1*, extending beyond a TAD boundary present in HAP1s, embryonic stem cells (ESCs), neural progenitor cells (NPCs) and neurons (Fig. 3). These contacts suggest that *FOXG1* regulation extends beyond this TAD boundary, substantiated by the presence of inter boundary looping interactions as well (Supplementary Fig. 7-8).

**Figure 3:**
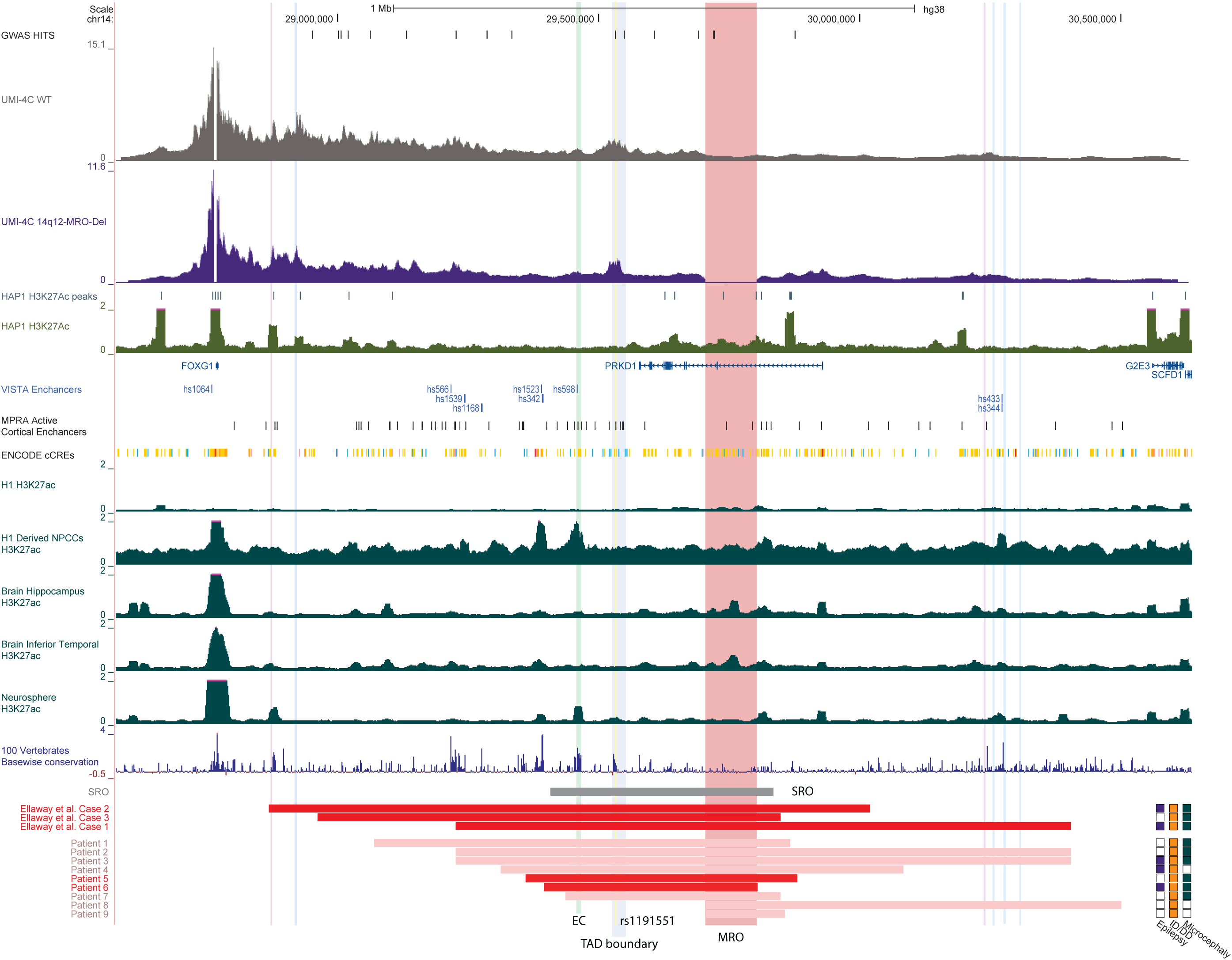
The *FOXG1* regulatory region is a complex landscape of multiple known and putative CREs encompassing patient-specific deletions. The MRO defined by the 14q12 patient deletions in this study is highlighted in red. *FOXG1* UMI-4C profiles represent genomic interactions of *FOXG1* in the 14q12 region (WT, grey; 14q12-MRO-Del; purple). ADHD- and schizophrenia-associated GWAS loci ^56^, active CREs from MPRAs in cerebral organoids and fetal brain samples ^33^, CREs defined by the enhancer histone modification, H3K27Ac, as well as VISTA enhancers and ENCODE Candidate CREs (cCREs) in this region overlap peaks from the UMI-4C contact profiles. Validated VISTA enhancers that show enhancer activity in transgenic mice are shown in blue, and region specificity shown in Supplementary Fig. 5 ^30,31^. A previously described *FOXG1* regulatory region, SRO (smallest region of overlap, horizontal grey bar) ^25^ in 14q12 encompasses VISTA enhancer hs598 with validated enhancer activity in the neural tube (Supplementary Fig. 5 and 31) as well as an enhancer cluster (EC) with validated enhancer activity in zebrafish development (mint vertical highlight) ^32^. The schizophrenia-associated SNP rs1191551 (yellow vertical highlight) ^57^ and the distal *FOXG1* TAD boundary (grey vertical highlight) also lie within the SRO. Abbreviations: NPCCs, Neural Progenitor Cultured Cells. Differential UMI contacts between WT and 14q12-MRO-Del are represented as pink (positive log2FoldChange) and blue (negative log2FoldChange) vertical highlights. ENCODE cCRE legend: Promoter-like signature (red), Proximal (orange) and Distal (yellow) enhancer-like signature, DNAse H3K4me3 (pink) and CTCF-only (blue).

As expected, in the 14q12-MRO-Del, physical genomic interactions were lost between the *FOXG1* promoter and the MRO locus (Fig. 3 and Fig. 4A). Deletion of the 14q12-MRO results in the deletion of at least two, but possibly more enhancers, as defined by H3K27Ac peaks in the HAP1 cell line (Fig. 4A). We selected these two H3K27Ac peaks (cCRE1 and cCRE2) in the MRO to test for enhancer activity in the HAP1 cell line and the HT-22 cell line derived from mouse hippocampus. cCRE1, which has high levels of conservation, showed a statistically significant 1.6-fold and 3.4-fold increase in expression in HAP1s and HT-22s, respectively (Fig. 4B). Conversely, cCRE2 reduced expression of the reporter by 2.5 - 4.8-fold in each cell type. Due to size constraints, we could not test both cCREs simultaneously, which would be more reflective of the context specific regulatory potential of the MRO. Nevertheless, in this system, our data suggests that both cCRE1 and cCRE2 have regulatory potential, and deletion of at least the cCRE1 element could underpin the reduced *FOXG1* expression elicited by MRO deletion. Given that both the cCRE H3K27Ac peaks are present in stem cell derived neural precursor lines and the human brain (Fig. 4A), and that there are two additional active regulatory elements (Fig. 4A - MPRA active 1/2,^33^) in the developing human brain within the MRO, we collectively conclude that the MRO locus is likely regulatory in a neuronal context and MRO loss results in reduced *FOXG1* expression.

**Figure 4:**
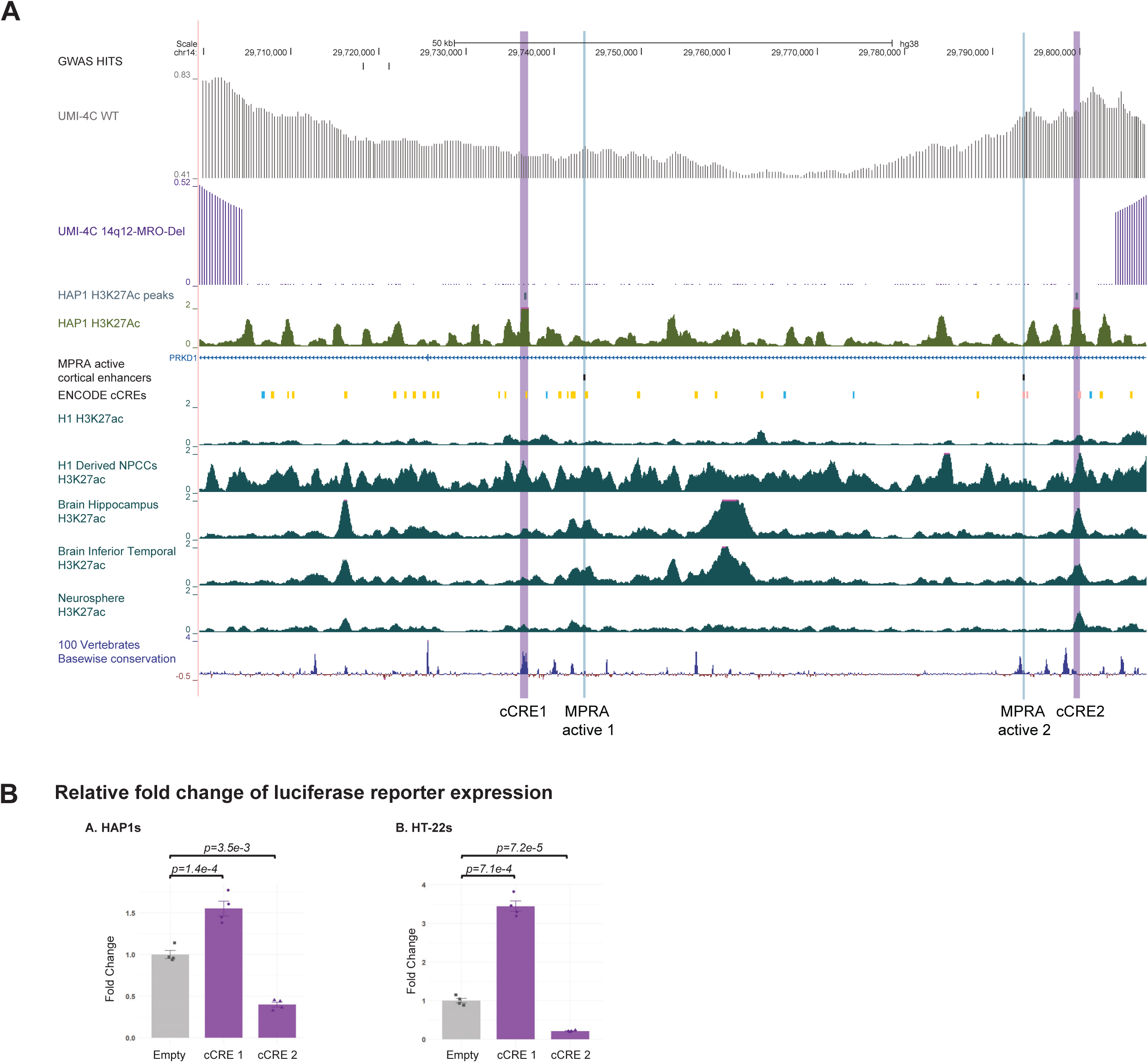
Evaluation of *FOXG1* CREs in the MRO. (A) A focus on the MRO region showcases native *FOXG1* genomic interactions in WT that are lost in 14q12-MRO-Del. Two active CREs (cyan highlights) from MPRAs in cerebral organoids and fetal brain samples reside within the MRO.^33^ Putative *FOXG1* CREs (cCRE1/2, purple highlights) in the MRO are identified by the two H3K27Ac peaks conserved in HAP1s (ENCODE) and neural cell types (RoadMap Epigenomic Project). (B) Dual luciferase assay in HAP1s (human) and HT-22s (mouse) shows luciferase reporter expression is increased in the presence of cCRE1 and decreased in the presence of cCRE2 (n =4 for Empty, cCRE1 and cCRE2). Abbreviations: NPCCs, Neural Progenitor Cultured Cells. ENCODE cCRE legend: Distal enhancer-like signature (yellow), DNAse H3K4me3 (pink) and CTCF-only (blue).

### Gene *expression* analysis identifies shared DEGs between 14q12-MRO-Del and *FOXG1*^KO^

Given *FOXG1’s* role as a transcription factor, we evaluated the effects of *FOXG1* downregulation on overall gene expression changes using RNA-seq. We identified 1036 differentially expressed genes (DEGs) in 14q12-MRO-Del, and 517 DEGs in FOXG1^KO^, compared to WT. *FOXG1* was a DEG in both lines with a larger effect in FOXG1^KO^ (log2FC −2.753916 and adjusted p-value 8.484237e-34), as compared to the 14q12-MRO-Del line (log2FC −0.7397485 and adjusted p-value 0.002290243). Furthermore, we identified 86 DEGs that overlapped between the two datasets. The shared DEGs included genes involved in cell cycle regulation (*CCND1, BMP7)* ^38–40^ as well as those implicated in epilepsy (*FOSL2, ELAVL3, EPHA5*, *OLFM3, ARHGAP4)* ^41–45^, ID/microcephaly (*XYLT1, COLEC11, CADPS2)* ^46–48^, and other NDDs (*KY, ADGRL3, TCEAL9)* ^49–51^ (Fig. 5 and Supplementary Data 1). Functional enrichment analysis of DEGs revealed a few shared molecular functions and biological pathways between 14q12-MRO-Del and FOXG1^KO^ including terms relevant in a neuronal context: “regulation of nervous system development”, “axon/neuron projection guidance” and “regulation of neurotransmitter levels” (Supplementary Fig. 9). We also used two existing chromatin immunoprecipitation sequencing experiments from mouse embryonic brain (E15.5 cortex ^52^ and E16.5 ganglionic eminence ^53^) to identify 8007 genes bound by Foxg1. Of these direct Foxg1 targets, a comparable fraction were also DEGs in the FOXG1^KO^ and 14q12-MRO-Del lines (17% (86/517 DEGs) in FOXG1^KO^ and 15% (154/1036 DEGs) in 14q12-MRO-Del) (Supplementary Data 1). These results are comparable to the 33% of DEGs that are direct targets of Foxg1 in the mouse ganglionic eminence ^53^. Thus collectively, even though we modeled MRO loss in a non-neuronal HAP1 cell line, we observe overlapping DEGs, including direct Foxg1 targets, in the MRO deletion and *FOXG1* coding loss, suggesting shared downstream effects of reduced expression of this transcription factor.

**Figure 5:**
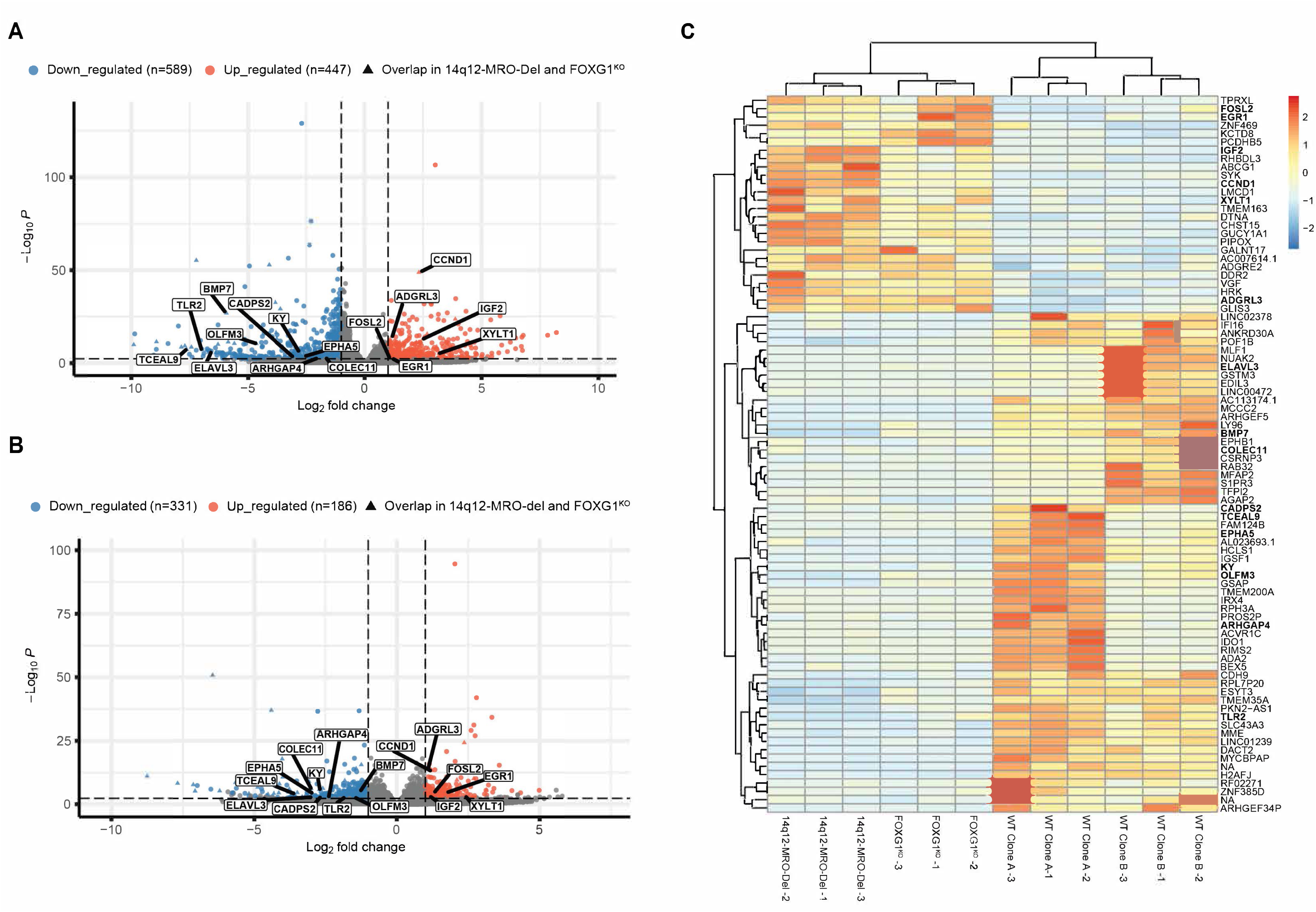
Differential gene expression analysis in MRO and *FOXG1* loss. Volcano plot of DEGs in (A) 14q12-MRO-Del vs WT and (B) FOXG1^KO^ vs WT highlighting down regulated (blue: log2FC<1) and upregulated (orange:log2FC>1) DEGs (adjusted p-value<0.05). Genes that overlap both MRO and FOXG1 loss datasets are indicated by triangles. (C) Heatmap of gene expression changes for overlapping DEGs between 14q12-MRO-Del, and FOXG1^KO^ HAP1s. (n=3 for WT Clone A, WT Clone B, 14q12-MRO-Del and FOXG1^KO^). Genes implicated in *FOXG1* pathophysiology and clinical phenotypes are highlighted (panels A and B) and appear in bold in panel C.

## Discussion

*FOXG1* is a dosage sensitive gene where both over and under expression are associated with epilepsy and NDDs ^16,54,55^. SVs, especially microdeletions in the 14q12 region downstream of *FOXG1* are reported in several individuals with clinical phenotypes that resemble *FOXG1* haploinsufficiency ^23,25–27^. Whether the observed phenotypes are a result of the deletions remains unclear, and the complex mechanisms of *FOXG1* regulation by CREs in this region, and how the microdeletions might disrupt this landscape are not fully understood. We identified nine 14q12 microdeletions downstream of *FOXG1* in DECIPHER in individuals with epilepsy, ID and/or microcephaly, and having no other clear causal variants. The deletions ranged from ∼151.8 kb to 1.17 Mb in size and were of mostly unspecified pathogenicity. We identified a shared 0.1 Mb region (MRO), deleted in all patients, nearly 0.93 Mb distal to *FOXG1* that regulates *FOXG1* expression.

Deletion of the MRO led to reduced *FOXG1* expression by roughly half in the HAP1 cell line, possibly due to the deletion of cCRE1 which activated expression of a reporter gene in HAP1s. This cCRE is highly conserved across evolution and correlates with a peak of H3K27Ac in HAP1s and a distal enhancer signature in ENCODE, collectively suggesting it functions as an enhancer in HAP1s. However, the second H3K27Ac peak, cCRE2 which is poorly conserved, reduced expression of the reporter gene in HAP1s and shows correlation with H3K4me3 and DNAse hypersensitivity from ENCODE CREs usually demarcating promoter regions. Taken together, the impact of cCRE2 is less clear; while there is good evidence that cCRE1 deletion may reduce expression of *FOXG1* in HAP1s. Further, cCRE1 increased expression of a reporter gene to an even greater extent in a mouse hippocampal cell line and also shows some evidence of H3K27Ac enrichment in a neuronal context. Moreover, there are two additional active enhancers in the MRO detected from primary cortical fetal brain tissue ^33^. Collectively, our data in HAP1s and HT-22s, as well as our reanalysis of published data, suggests that the MRO has the potential to regulate *FOXG1* expression, and that deletion of this region leads to reduced FOXG1 and contributes to the development of neurological features in individuals with 14q12 deletions.

Deletion of the MRO reduced *FOXG1* expression but did not eliminate it, indicating that multiple CREs across the region need to be deleted to abrogate *FOXG1* expression. Indeed, the MRO lies within a larger regulatory region of *FOXG1* designated as the SRO (smallest region of overlap); previously defined by *de novo* 14q12 deletions downstream of *FOXG1* in three individuals with *FOXG1* syndrome related phenotypes including epilepsy, ID, developmental delays and severe postnatal microcephaly. The SRO is a complex region with multiple validated, highly conserved active enhancers in mouse, zebrafish and human fetal brain tissue and organoid models ^30–33^. We refined the SRO to the MRO based on the distal deletions in patient 8 and 9. Of note, these two individuals present with ID only. In contrast, individuals with deletion of both the SRO and MRO (patient 1-7) generally present with more severe clinical features of *FOXG1* haploinsufficiency: ID, seizures and microcephaly (**Table 1**). These varied clinical presentations, taken together with the reduction, but not complete loss of *FOXG1* by MRO deletion, suggests that disruption of multiple CREs is required to completely ablate *FOXG1* expression. Moreover, the number of disrupted CREs may underpin genotype-phenotype relationships; where individuals with deletion of more CREs have lower *FOXG1* expression and clinical features more reminiscent of individuals with *FOXG1* coding variants, while individuals with fewer CREs deleted (e.g. patients 8 and 9) have less severe clinical presentation correlated with higher FOXG1 abundance. Indeed, this modulated view of *FOXG1* expression finds support in other neurological conditions. For instance, there are multiple GWAS loci in the 14q12 regulatory region that have been associated with increased risk for ADHD and schizophrenia ^56^. Deletion of a ∼500 bp region flanking a schizophrenia-associated SNP rs1191551 reduced *FOXG1* expression by ∼20% in HEK293T cells ^57^. Moreover, a luciferase assay demonstrated reduced activity of the risk rs1191551 allele, suggesting that even this single nucleotide change can modulate *FOXG1* expression. Collectively, these studies highlight the need to systematically assess both disease-associated SVs and single nucleotide changes across this regulatory region in neurological disease broadly, and that variable gene expression across neurodevelopment may have varied phenotypic consequences. As genome sequencing facilitates more widespread and smaller SV and SNV detection, these genotype-phenotype relationships may become more apparent.

In addition to the loss of contacts in the MRO, we also observed increased distal contacts beyond a putative TAD boundary in the 14q12-MRO-Del HAP1 line. This included three increased contacts with regions that harbor forebrain-specific, active VISTA enhancers, and are supported by contact maps in human NPCs and neurons as well. These results support the notion of enhancer redundancy across the 14q12 locus. Tight regulation via multiple CREs is a known characteristic of developmental genes, particularly those that are exquisitely sensitive to dosage during mammalian development ^58^. This redundancy is likely a protective mechanism against deleterious genomic aberrations ^59^. For instance, individual deletion of each of ten regulatory elements involved in mouse limb development resulted in normal limb phenotypes, whereas pairwise deletions of these loci resulted in noticeably different limb formations, demonstrating partial redundancy ^59^. These redundant, or shadow enhancers regulate the same target gene and overlap in part or full, in the developmental space and time to fine tune gene dosage ^58^. Similarly, *FOXG1* expression is subject to tight spatiotemporal regulation during corticogenesis. Furthermore, the effect of shadow enhancers on gene expression is not always additive; depending on the availability of transcription factors at each of the enhancers and the buffering requirement of the cell, the effects can be superadditive, subadditive, or even repressive ^58,60^. Here we demonstrate that there are multiple CREs within the MRO that may regulate *FOXG1* expression, but that there are likely additional redundant enhancers across the 14q12 locus region. Testing this hypothesis, however, is challenging, and a limitation of the dual sgRNA guide genome-editing strategy used in this study to introduce the ∼100 kb MRO deletion. While newer methods such as dCas9-controlled CRISPR/Cas3 can generate deletions on a megabase scale ^61^, our approach has too low an efficiency to introduce the >400 kb SRO. Future studies, using Cas3-like deletion approaches in cellular and animal models are key for examining the redundancy of CREs in the 14q12 region, and the absolute number of CREs that need to be deleted to abrogate *FOXG1* expression. Furthermore, massively parallel reporter assays (MPRA) in human neurodevelopmental stem cell models and human tissues, such as those described ^33^, are likely to be essential for assessing the activity of individual CREs.

We examined the transcriptional profile of the MRO deletion and FOXG1 knockout lines, demonstrating overlap in the genes that were differentially expressed, notably BMP family protein *BMP7* ^40^, *CCND1*, a G1 phase cyclin ^39^ and *EGR1*, a transcription factor and member of the EGR-*FOXG1* coupled regulatory network ^62^. Of note, these genes have overlap with the known roles of FOXG1 in a neuronal context, including modulating BMP signaling and the cell cycle, ensuring cells remain in proliferative states ^19,40^. As well as these single genes, key biological pathways of neurodevelopment were also enriched for DEGs, including overlapping genes in both models of *FOXG1* reduction. While these results are encouraging, they do highlight a limitation of our study, in that we do not demonstrate the impact of MRO loss in a neuronal cell line. We selected HAP1s because of the ease of editing and robust FOXG1 abundance, and are confident that at least some of the regulatory landscape is overlapping, but we acknowledge that this is not a neuronal line. We did also generate iPSC lines heterozygous for MRO deletion and differentiated them to neural cell types. However, we found inconsistent *FOXG1* expression in iPSC-derived NPCs and iPSC-derived neurons that was not reproducible, even in wildtype control lines (data not shown). High variability in the endogenous expression of this gene is also seen in previous iPSC studies ^20,40^. Therefore, collectively we could not use these neuronal models as an effective experimental system to study the effects of MRO- and *FOXG1* loss. In the future, reporter-based systems, and live-imaging will be necessary to accurately quantify the temporal expression of FOXG1 in iPSC-derived models, to counteract the variability inherent in these systems.

In summary, our work refines the regulatory landscape of the *FOXG1* locus, and we identify a ∼0.1 Mb regulatory region downstream of *FOXG1* in the 14q12 region. Within this MRO, we define putative *FOXG1* CREs and emphasize that this locus exhibits both enhancer redundancy as well as shadow enhancer activity. The complex regulatory nature of the *FOXG1* locus is reflected in its importance in disease etiology. We define the causal mechanisms of a subset of clinical features of *FOXG1* syndrome and posit that deletion of multiple CREs is necessary to cause *FOXG1* haploinsufficiency. Overall, this critical regulatory locus should be examined across both NDDs and neuropsychiatric disease for causative and contributory variants, presenting unique opportunities for gene-targeting therapies via modulation of *FOXG1* expression.

## Methods

### Cell line maintenance

Human eHAP (fully haploid engineered HAP1) cells from Horizon Discovery BioSciences (C669) were maintained in IMDM (Thermo Fisher Scientific) supplemented with 10% Tet-approved FBS (Atlanta Biologicals) and 1% penicillin/streptomycin (Thermo Fisher Scientific). Mouse hippocampal HT-22 cells (Sigma SCC129) were maintained in DMEM containing 4.5 g/l glucose (Thermo Fisher Scientific) supplemented with 10% Tet-approved FBS (Atlanta Biologicals) and 1% penicillin/streptomycin (Thermo Fisher Scientific). The cells were tested for mycoplasma contamination, and all experiments were performed within 3 to 10 passages.

### Selection of candidate 14q12 microdeletions and cell line generation

We used DECIPHER (DatabasE of genomiC varIation and Phenotype in Humans using Ensembl Resources) ^63^ to identify 14q12 microdeletions downstream of the NDD-related gene, *FOXG1* in individuals with epilepsy and related NDDs. Deletions within *FOXG1’s* TAD that excluded *FOXG1* and extended up to hg38: chr14:30,550,000 (approaching the distal TAD boundary limit) were considered ^64^. Larger deletions spanning multiple genes were excluded as gene dosage effects from multiple encompassed genes are likely to confound patient phenotypes.

CRISPR genome editing was used to create 14q12-MRO-Del and FOXG1^KO^ HAP1 lines. Guide RNAs were designed to target each of the two breakpoints of the MRO and cloned into a Cas9-GFP vector (PX458, Addgene #48138) or a Cas9-mCherry vector (a PX458 derivative engineered in-house) (Supplementary Table 1 for guide RNA sequences). Dual guide electroporation was performed using the Neon Transfection System. 24-48h post transfection, FACS was performed for single cell clonal isolation and expansion of GFP and mCherry double positive cells. Clonal lines were genotyped by PCR, followed by Sanger sequencing, to detect the presence of the deletion (Supplementary Table 1 for primer sequences). Whole genome sequencing (WGS) was performed to further confirm the intended genotype. Moreover, we used the MetaSV pipeline to detect the presence of any large unintended SVs, and any lines harboring such variants were excluded ^65^. FOXG1^KO^ lines were similarly generated, with dual guide RNAs targeting the start and end of *FOXG1*’s coding region (Supplementary Table 1). Clones of 14q12-MRO-Del and FOXG1^KO^, along with the parental/unedited line (WT Clone A) and a wildtype clone from the CRISPR screen (WT Clone B) as isogenic controls, were selected for experimentation (Supplementary Table 2).

### *FOXG1* mRNA and protein quantification

Quantitative reverse transcription PCR was performed to assay *FOXG1* mRNA levels. RNA was extracted and purified from HAP1s in culture using TRIzol reagent (Invitrogen #15596026). qRT-PCR reactions were performed using Biorad iTaq Universal One-Step RT-qPCR Kit (1725150). 25ng RNA per reaction was used and triplicate qRT-PCR reactions were carried out for each target (Supplementary Table 1) on a Biorad CFX384 Real-Time PCR Detection system. Raw Ct values were obtained for each sample. Ct values for *FOXG1* were transformed by first normalizing to EAR and then normalizing to WT to calculate relative fold change of *FOXG1* ^66^. Two Sample T-Test (Welch’s T-Test) was performed to test statistical significance between samples. *PRKD1* mRNA quantification was similarly performed.

Western blotting was performed to assay FOXG1 protein levels. Whole cell lysates were prepared using RIPA Lysis and Extraction Reagent (Thermo Scientific 89900) in the presence of protease inhibitors (cOmplete™, Mini, EDTA-free Protease Inhibitor Cocktail, Sigma 11836170001). 20 μg of protein per well was separated on NuPage 3-8% Tris-Acetate precast gels (Invitrogen) and transferred onto a PVDF membrane (Millipore). Immunoblotting was performed using primary antibodies (Supplementary Table 3) and detected with peroxidase-conjugated secondary antibodies (Supplementary Table 3). This was followed by detection with ECL peroxidase substrate (Cytiva Amersham RPN2232). Protein band intensities on immunoblots were quantified by densitometric analysis. Two Sample T-Test (Welch’s T-Test) was performed to test statistical significance between samples.

### Locus specific chromatin capture

UMI-4C was performed for targeted and quantitative capture of *FOXG1*’s genomic interactions with the following specifications ^14^. *FOXG1* viewpoint was defined as a 1 kb window at the 5’ end of the gene. Crosslinked chromatin from 5 x 10^6^ cells were used per sample. UMI-4C libraries were constructed by enriching the 4C library for *FOXG1* contacts using viewpoint specific primers (Supplementary Table 1) ^32^. Two unique barcode adapters per sample were utilized to generate five UMI-4C reactions per adapter, which were pooled to make a final UMI-4C library. UMI-4C experiments were performed at least twice per clone. Libraries were subject to 75 bp paired end sequencing on the Illumina MiniSeq platform at a sequencing depth of 1-2M reads per sample.

Raw read quality was assessed using FastQC ^67^. The fastq files from sequencing were analyzed using the UMI-4C R Package as described ^14,68^. Since the UMI-4C library is made up of adapter ligated DNA fragments that consist of both the bait and its spatially proximal genomic region, it generates sequencing reads that map to both these loci. To quantify these spatially proximal genomic interactions, the R package transforms the raw sequencing reads to unique genomic interactions (UMI counts) per genomic restriction fragment and the resulting data is saved as genomic tracks. From these tracks, we extracted viewpoint-specific UMI counts for a given region of interest using the package. UMI counts were normalized to the total UMI counts within the region of interest and a smoothed interaction profile was produced using default parameters in the package. In this way, we generated UMI-4C profiles for the 14q12 region of interest (hg38: chr14: 28,965,045-30,631,045) for WT and 14q12-MRO-Del samples. Differential genomic interactions between samples were evaluated using the UMI4Cats R package ^69^. Significant differential contacts between WT and 14q12-MRO-Del were inferred by the Wald Test using the waldUMI4C() function with FDR adjusted *p-*value of 0.05.

### Analysis of H3K27Ac data across 14q12 *FOXG1* regulatory region

H3K27Ac ChIP-seq data was assessed from published data in HAP1s (GSE167721), adult brain hippocampus (GSM1112791) and temporal lobe (GSM772995), the embryonic stem cell line H1 (GSM466732) as well as neural precursor cells (NPCs) derived from H1 (GSM753429) and ganglionic eminence neurospheres derived from H1 (GSM1127083).

### Validation of MRO cCREs using luciferase assay

The regulatory activity of cCREs was evaluated in HAP1 and HT-22 cell lines using the Dual-Luciferase® Reporter Assay System (Promega E1910). cCRE sequences (Supplementary Table 5) were cloned into the pGL4.23[luc2/minP] vector (Promega E8411) containing firefly luciferase. The pGL4.74[hRluc2TK] Vector (Promega E6921) with *Renilla* luciferase was used as internal transfection control. pGL4.23 minus an upstream cCRE sequence (Empty) was used as control for baseline firefly luciferase expression. Dual transfection of pGL4.23 and pGL4.74 plasmids was performed using Xfect (Takara Bio) for HAP1s, and Lipofectamine (Thermo Fisher Scientific) for HT-22s. Cells were assayed 48h post transfection. Luminescence from Firefly luciferase was first normalized to the internal *Renilla* luciferase control and then further normalized to the Empty control to calculate relative fold change of luciferase reporter expression with cCREs. Experiments were repeated four times.

### Bulk RNA-Seq library prep and data analysis

RNA was extracted and purified from HAP1s in culture by the guanidinium thiocyanate-phenol-chloroform extraction method using TRIzol^TM^ reagent (Invitrogen #15596-026). Purified RNA samples were assayed on a TapeStation 4150 (Agilent) to determine the concentration and RIN (RNA Integrity Number) values. RNA samples with RIN > 9 were sent to Novogene for mRNA library preparation and subject to 150 bp paired end sequencing on a NovaSeq platform at a sequencing depth of at least 20 M reads per sample.

Raw read quality was assessed using FastQC ^67^. Reads were trimmed to remove adapter sequences using HTStream, aligned to the human reference genome GRCh38/hg38 and GENCODE v38 using STAR ^70^. The read count matrix was generated using GENCODE v38 using Rsubread ^71^. Differentially expressed genes (DEGs) were detected using DESeq2 using an adjusted p-value threshold of 0.05 and log2Fold change greater/less than absolute value of 1 ^72^. Functional enrichment analysis was performed using gene ontology in the clusterProfiler package using a Benjamini-Hochberg corrected p-value threshold of 0.05 ^73–76^. Published FOXG1 peaks were retrieved from prior work ^52,53^ and the overlapping (i.e. present in either dataset) set of genes (n=8007) were used to identify the direct FOXG1 genes that were also DEGs.

### Reporting summary

Further information on research design is available in the Nature Research Reporting Summary linked to this article.

## Data availability

The main data supporting the findings of this study are available within the article and its Supplementary Information. Source data are provided with this paper. UMI-4C and RNA-Seq raw data files from this study are available at the NCBI SRA database with BioProject accession number PRJNA1348521. Additional details on datasets and protocols of this study will be made available by the corresponding author upon reasonable request.

## Author contributions

AR, MB, LA, OJ, EY, JZC and CB generated the experimental data. NB and JDC assisted with experiments. AR, JDC, GLC performed data analysis and interpretation of results. GLC and AR conceived the presented idea, developed the theory, wrote this manuscript and generated figures for this manuscript. GLC supervised the project.

## Acknowledgements

Our work was supported by the following funding sources: the Chicago Biomedical Consortium Catalyst Award to GLC and the American Epilepsy Society Predoctoral Fellowship to AR.

## References

1. McRae, J. F. et al. Prevalence and architecture of de novo mutations in developmental disorders. Nature 2017 542:7642 542, 433–438 (2017).

2. Jansen, S., Vissers, L. E. L. M. & de Vries, B. B. A. The Genetics of Intellectual Disability. Brain Sciences 2023*, Vol.* 13, *Page 231* 13, 231 (2023).

3. Kaminsky, E. B. et al. An evidence-based approach to establish the functional and clinical significance of copy number variants in intellectual and developmental disabilities. Genet Med 13, 777–784 (2011).

4. D’haene, E. & Vergult, S. Interpreting the impact of noncoding structural variation in neurodevelopmental disorders. Genetics in Medicine 23, 34–46 (2021).

5. Hebbar, M. & Mefford, H. C. Recent advances in epilepsy genomics and genetic testing. F1000Res 9, (2020).

6. Helbig, K. L. et al. Diagnostic exome sequencing provides a molecular diagnosis for a significant proportion of patients with epilepsy. Genet Med 18, 898–905 (2016).

7. Retterer, K. et al. Clinical application of whole-exome sequencing across clinical indications. Genet Med 18, 696–704 (2016).

8. Cauwelier, B. et al. Molecular cytogenetic study of 126 unselected T-ALL cases reveals high incidence of TCRβ locus rearrangements and putative new T-cell oncogenes. Leukemia 2006 20:*7* 20, 1238–1244 (2006).

9. Weischenfeldt, J. et al. Pan-cancer analysis of somatic copy-number alterations implicates IRS4 and IGF2 in enhancer hijacking. Nature Genetics 2016 49:*1* 49, 65–74 (2016).

10. Wang, X. et al. Genome-wide detection of enhancer-hijacking events from chromatin interaction data in rearranged genomes. Nature Methods 2021 18:*6* 18, 661–668 (2021).

11. Tenney, A. P. et al. Noncoding variants alter GATA2 expression in rhombomere 4 motor neurons and cause dominant hereditary congenital facial paresis. Nature Genetics 2023 55:*7* 55, 1149–1163 (2023).

12. Ibn-Salem, J. et al. Deletions of chromosomal regulatory boundaries are associated with congenital disease. Genome Biol 15, 423 (2014).

13. Lupiáñez, D. G. et al. Disruptions of Topological Chromatin Domains Cause Pathogenic Rewiring of Gene-Enhancer Interactions. Cell 161, 1012–1025 (2015).

14. Mohajeri, K. et al. Transcriptional and functional consequences of alterations to MEF2C and its topological organization in neuronal models. Am J Hum Genet 109, 2049–2067 (2022).

15. Redin, C. et al. The genomic landscape of balanced cytogenetic abnormalities associated with human congenital anomalies. Nat Genet 49, 36–45 (2017).

16. Hettige, N. C. & Ernst, C. FOXG1 Dose in Brain Development. Front Pediatr 7, 482 (2019).

17. Xuan, S. et al. Winged helix transcription factor BF-1 is essential for the development of the cerebral hemispheres. Neuron 14, 1141–1152 (1995).

18. Dastidar, S. G., Landrieu, P. M. Z. & D’Mello, S. R. FoxG1 Promotes the Survival of Postmitotic Neurons. Journal of Neuroscience 31, 402–413 (2011).

19. Hanashima, C., Shen, L., Li, S. C. & Lai, E. Brain Factor-1 Controls the Proliferation and Differentiation of Neocortical Progenitor Cells through Independent Mechanisms. Journal of Neuroscience 22, 6526–6536 (2002).

20. Hettige, N. C. et al. FOXG1 dose tunes cell proliferation dynamics in human forebrain progenitor cells. Stem Cell Reports 17, 475–488 (2022).

21. Ariani, F. et al. FOXG1 Is Responsible for the Congenital Variant of Rett Syndrome. The American Journal of Human Genetics 83, 89–93 (2008).

22. Mitter, D. et al. FOXG1 syndrome: genotype–phenotype association in 83 patients with FOXG1 variants. Genetics in Medicine 2018 20:*1* 20, 98–108 (2017).

23. Kortüm, F. et al. The core FOXG1 syndrome phenotype consists of postnatal microcephaly, severe mental retardation, absent language, dyskinesia, and corpus callosum hypogenesis. J Med Genet 48, 396–406 (2011).

24. Brunetti-Pierri, N. et al. Duplications of FOXG1 in 14q12 are associated with developmental epilepsy, mental retardation, and severe speech impairment. European Journal of Human Genetics 2011 19:*1* 19, 102–107 (2010).

25. Allou, L. et al. 14q12 and severe Rett-like phenotypes: new clinical insights and physical mapping of FOXG1-regulatory elements. Eur J Hum Genet 20, 1216–1223 (2012).

26. Ellaway, C. J. et al. 14q12 microdeletions excluding FOXG1 give rise to a congenital variant Rett syndrome-like phenotype. European Journal of Human Genetics 2013 21:*5* 21, 522–527 (2012).

27. Mehrjouy, M. M. et al. Regulatory variants of FOXG1 in the context of its topological domain organisation. European Journal of Human Genetics 2017 26:*2* 26, 186–196 (2017).

28. Karczewski, K. J. et al. The mutational constraint spectrum quantified from variation in 141,456 humans. Nature 2020 581:7809 581, 434–443 (2020).

29. Rausell, A. et al. Common homozygosity for predicted loss-of-function variants reveals both redundant and advantageous effects of dispensable human genes. Proc Natl Acad Sci U S A 117, 13626–13636 (2020).

30. Visel, A., Minovitsky, S., Dubchak, I. & Pennacchio, L. A. VISTA Enhancer Browser--a database of tissue-specific human enhancers. Nucleic Acids Res 35, (2007).

31. Kosicki, M. et al. VISTA Enhancer browser: an updated database of tissue-specific developmental enhancers. Nucleic Acids Res 53, (2025).

32. Hamerlinck, L. et al. Non-coding structural variants identify a commonly affected regulatory region steering FOXG1 transcription in early neurodevelopment. medRxiv 2025.03.10.25323301 (2025) doi:10.1101/2025.03.10.25323301.

33. Deng, C. et al. Massively parallel characterization of regulatory elements in the developing human cortex. Science (1979) 384, 868–876 (2024).

34. Dunham, I. et al. An integrated encyclopedia of DNA elements in the human genome. Nature 489, 57–74 (2012).

35. Abascal, F. et al. Expanded encyclopaedias of DNA elements in the human and mouse genomes. Nature 2020 583:7818 583, 699–710 (2020).

36. Hitz, B. C. et al. The ENCODE Uniform Analysis Pipelines. bioRxiv (2023) doi:10.1101/2023.04.04.535623.

37. Luo, Y. et al. New developments on the Encyclopedia of DNA Elements (ENCODE) data portal. Nucleic Acids Res 48, D882–D889 (2020).

38. Pedraza, N. et al. Cytoplasmic expression of the cell cycle regulator cyclin D1 in radial glial progenitor cells modulates brain cortex development. bioRxiv 2024.12.23.630056 (2024) doi:10.1101/2024.12.23.630056.

39. Lottini, G. et al. Zika virus induces FOXG1 nuclear displacement and downregulation in human neural progenitors. Stem Cell Reports 17, 1683 (2022).

40. Hettige, N. C. et al. FOXG1 targets BMP repressors and cell cycle inhibitors in human neural progenitor cells. Hum Mol Genet 32, 2511 (2023).

41. Cospain, A. et al. FOSL2 truncating variants in the last exon cause a neurodevelopmental disorder with scalp and enamel defects. Genetics in Medicine 24, 2475–2486 (2022).

42. Wutikeli, H., Xie, T., Xiong, W. & Shen, Y. ELAV/Hu RNA-binding protein family: key regulators in neurological disorders, cancer, and other diseases. RNA Biol 22, 1–11 (2025).

43. Liu, T. T. et al. [Changes in the expression of EphA5/ephrinA5 in the CA3 region of the hippocampus in rats with epilepsy and their role in the pathogenesis of temporal lobe epilepsy]. Zhongguo Dang Dai Er Ke Za Zhi 19, 1272–1277 (2017).

44. Tang, S. et al. Olfactomedin-3 Enhances Seizure Activity by Interacting With AMPA Receptors in Epilepsy Models. Front Cell Dev Biol 8, 557009 (2020).

45. Hu, Y. Y. et al. ARHGAP4 variants are associated with X-linked early-onset temporal lobe epilepsy. World J Pediatr 20, (2024).

46. LaCroix, A. J. et al. GGC Repeat Expansion and Exon 1 Methylation of XYLT1 Is a Common Pathogenic Variant in Baratela-Scott Syndrome. Am J Hum Genet 104, 35–44 (2019).

47. Gilissen, C. et al. Genome sequencing identifies major causes of severe intellectual disability. Nature 2014 511:7509 511, 344–347 (2014).

48. Bonora, E. et al. Maternally inherited genetic variants of CADPS2 are present in autism spectrum disorders and intellectual disability patients. EMBO Mol Med 6, 795–809 (2014).

49. Arif, B. et al. A novel homozygous KY variant causing a complex neurological disorder. Eur J Med Genet 63, 104031 (2020).

50. Menzel, M. H. M. M., Kohler, A., Hudemann, M., Jüngling, J. & Biskup, S. Case Report of a Juvenile Patient with Autism Spectrum Disorder with a Novel Combination of Copy Number Variants in ADGRL3 (LPHN3) and Two Pseudogenes. Application of Clinical Genetics 15, 125–131 (2022).

51. Albuainain, F. et al. Confirmation and expansion of the phenotype of the TCEAL1-related neurodevelopmental disorder. European Journal of Human Genetics 2024 32:*3* 32, 350–356 (2024).

52. Cargnin, F. et al. FOXG1 Orchestrates Neocortical Organization and Cortico-Cortical Connections. Neuron 100, 1083–1096.e5 (2018).

53. Sun, L. et al. Transcriptomic insights into fate choice of pallial versus subpallial GABAergic neurons. Nat Commun 16, (2025).

54. Li, J., Chang, H.. W., Lai, E., Parker, E.. J. & Vogt, P.. K. The oncogene qin codes for a transcriptional repressor - PubMed. Cancer Res 55, 5540–5544 (1995).

55. Seltzer, L. E. et al. Epilepsy and outcome in FOXG1-related disorders. Epilepsia 55, 1292–1300 (2014).

56. Cerezo, M. et al. The NHGRI-EBI GWAS Catalog: standards for reusability, sustainability and diversity. Nucleic Acids Res 53, D998–D1005 (2025).

57. Won, H. et al. Chromosome conformation elucidates regulatory relationships in developing human brain. Nature 538, 523 (2016).

58. Kvon, E. Z., Waymack, R., Gad, M. & Wunderlich, Z. Enhancer redundancy in development and disease. Nature Reviews Genetics 2021 22:*5* 22, 324–336 (2021).

59. Osterwalder, M. et al. Enhancer redundancy provides phenotypic robustness in mammalian development. Nature 2018 554:7691 554, 239–243 (2018).

60. Waymack, R., Fletcher, A., Enciso, G. & Wunderlich, Z. Shadow enhancers can suppress input transcription factor noise through distinct regulatory logic. Elife 9, 1–57 (2020).

61. Li, J. et al. Precise large-fragment deletions in mammalian cells and mice generated by dCas9-controlled CRISPR/Cas3. Sci Adv 10, 8052 (2024).

62. Hou, P. S., Miyoshi, G. & Hanashima, C. Sensory cortex wiring requires preselection of short- and long-range projection neurons through an Egr-Foxg1-COUP-TFI network. Nature Communications 2019 10:*1* 10, 1–18 (2019).

63. Firth, H. V. et al. DECIPHER: Database of Chromosomal Imbalance and Phenotype in Humans Using Ensembl Resources. The American Journal of Human Genetics 84, 524–533 (2009).

64. Wang, Y. et al. The 3D Genome Browser: A web-based browser for visualizing 3D genome organization and long-range chromatin interactions. Genome Biol 19, 1–12 (2018).

65. Mohiyuddin, M. et al. MetaSV: an accurate and integrative structural-variant caller for next generation sequencing. Bioinformatics 31, 2741–2744 (2015).

66. Livak, K. J. & Schmittgen, T. D. Analysis of relative gene expression data using real-time quantitative PCR and the 2(-Delta Delta C(T)) Method. Methods 25, 402–408 (2001).

67. Andrews, S. FastQC: A Quality Control Tool for High Throughput Sequence Data [Online]. Available online at: http://www.bioinformatics.babraham.ac.uk/projects/fastqc/. Preprint at https://qubeshub.org/resources/fastqc (2010).

68. Schwartzman, O. et al. UMI-4C for quantitative and targeted chromosomal contact profiling. Nat Methods 13, 685–691 (2016).

69. Ramos-Rodríguez, M., Subirana-Granés, M. & Pasquali, L. UMI4Cats: an R package to analyze chromatin contact profiles obtained by UMI-4C. Bioinformatics 37, 4240–4242 (2021).

70. Dobin, A. & Gingeras, T. R. Mapping RNA-seq Reads with STAR. Curr Protoc Bioinformatics 51, 11.14.1–11.14.19 (2015).

71. Liao, Y., Smyth, G. K. & Shi, W. The R package Rsubread is easier, faster, cheaper and better for alignment and quantification of RNA sequencing reads. Nucleic Acids Res 47, e47–e47 (2019).

72. Love, M. I., Huber, W. & Anders, S. Moderated estimation of fold change and dispersion for RNA-seq data with DESeq2. Genome Biol 15, 550 (2014).

73. Yu, G., Wang, L. G., Han, Y. & He, Q. Y. ClusterProfiler: An R package for comparing biological themes among gene clusters. OMICS 16, 284–287 (2012).

74. Wu, T. et al. clusterProfiler 4.0: A universal enrichment tool for interpreting omics data. The Innovation 2, 100141 (2021).

75. Yu, G. Thirteen years of clusterProfiler. The Innovation 5, 100722 (2024).

76. Xu, S. et al. Using clusterProfiler to characterize multiomics data. Nature Protocols 2024 19:*11* 19, 3292–3320 (2024).

