## Supplementary Information for "Analysis of 14q12 microdeletions reveals novel regulatory loci for the neurodevelopmental disorder-related gene, *FOXG1*"

**Ramamurthy et al.**

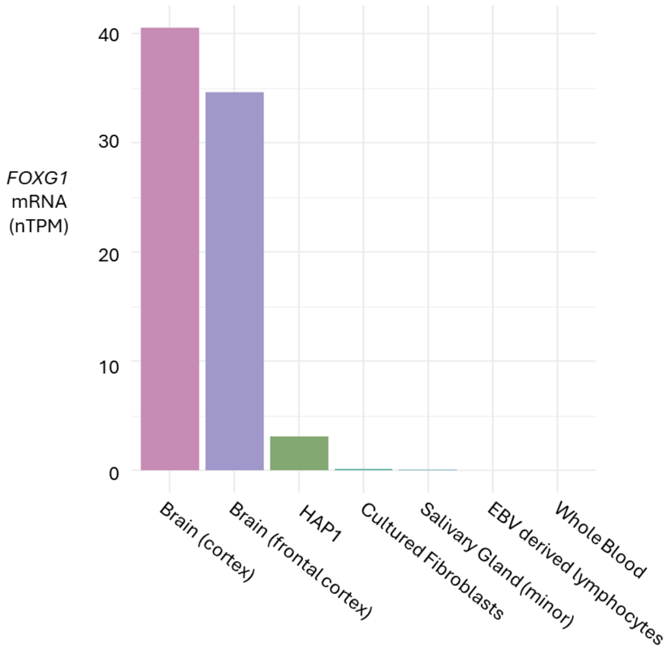


**Supplementary Figure 1.** *FOXG1* expression data. Normalized transcript per million (nTPM) values for *FOXG1* RNA were derived from GTEx for bulk tissues (GTEx 09.09.25, dbGaP accession number phs000424.vN.pN) and The Human Protein Atlas for HAP1 cell line (v24.proteinatlas.org).

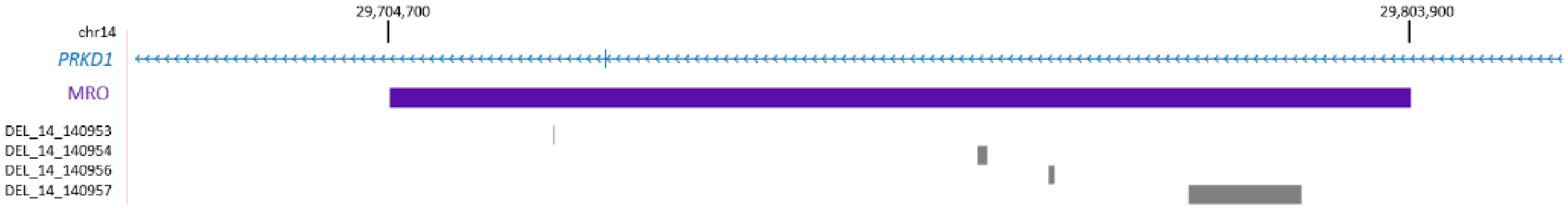


| Variant ID | Allele Count | Allele Frequency |
|---|---|---|
| DEL_14_140953 | 1 | 0.00006 |
| DEL_14_140954 | 2109 | 0.126424 |
| DEL_14_140956 | 1 | 0.00006 |
| DEL_14_140957 | 1 | 0.00006 |

**Supplementary Figure 2.** gnomAD variants, in particular deletions (grey bars), are mapped in the 14q12 MRO region (purple bar). The table reports allele count and minor allele frequency for each of these gnomAD deletions.

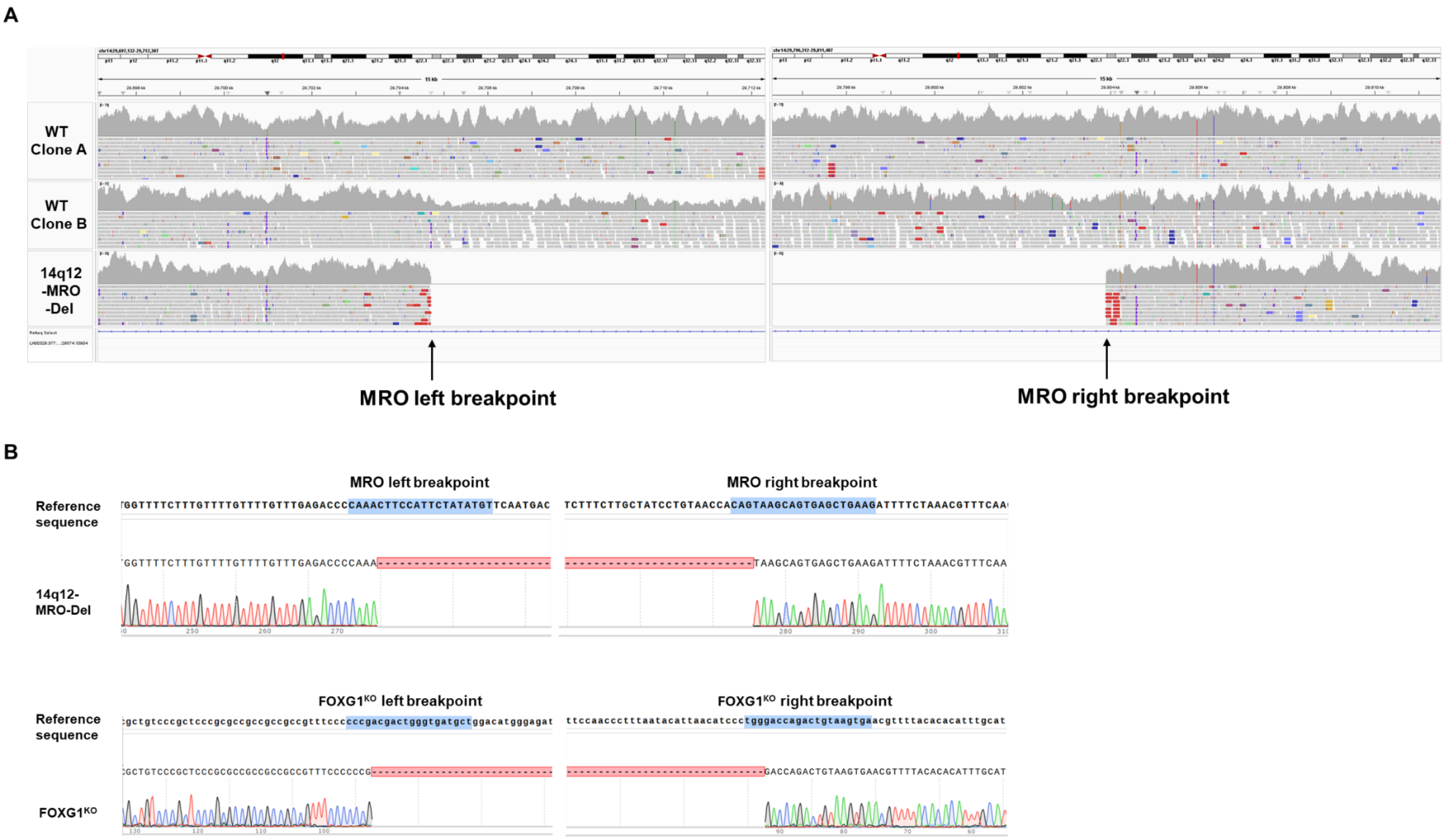


**Supplementary Figure 3**. Genotyping assays for validating CRISPR-Cas9 edited lines in this study. (A) Genome Sequencing data visualized using the Integrative Genomics Viewer confirm the deletion of MRO locus in 14q12-MRO-Del. (B) Deletion of the MRO in 14q12-MRO-Del and *FOXG1* coding sequence in FOXG1$^{KO}$ verified by Sanger sequencing. The guide sequence at each breakpoint is highlighted in blue within the reference sequence.

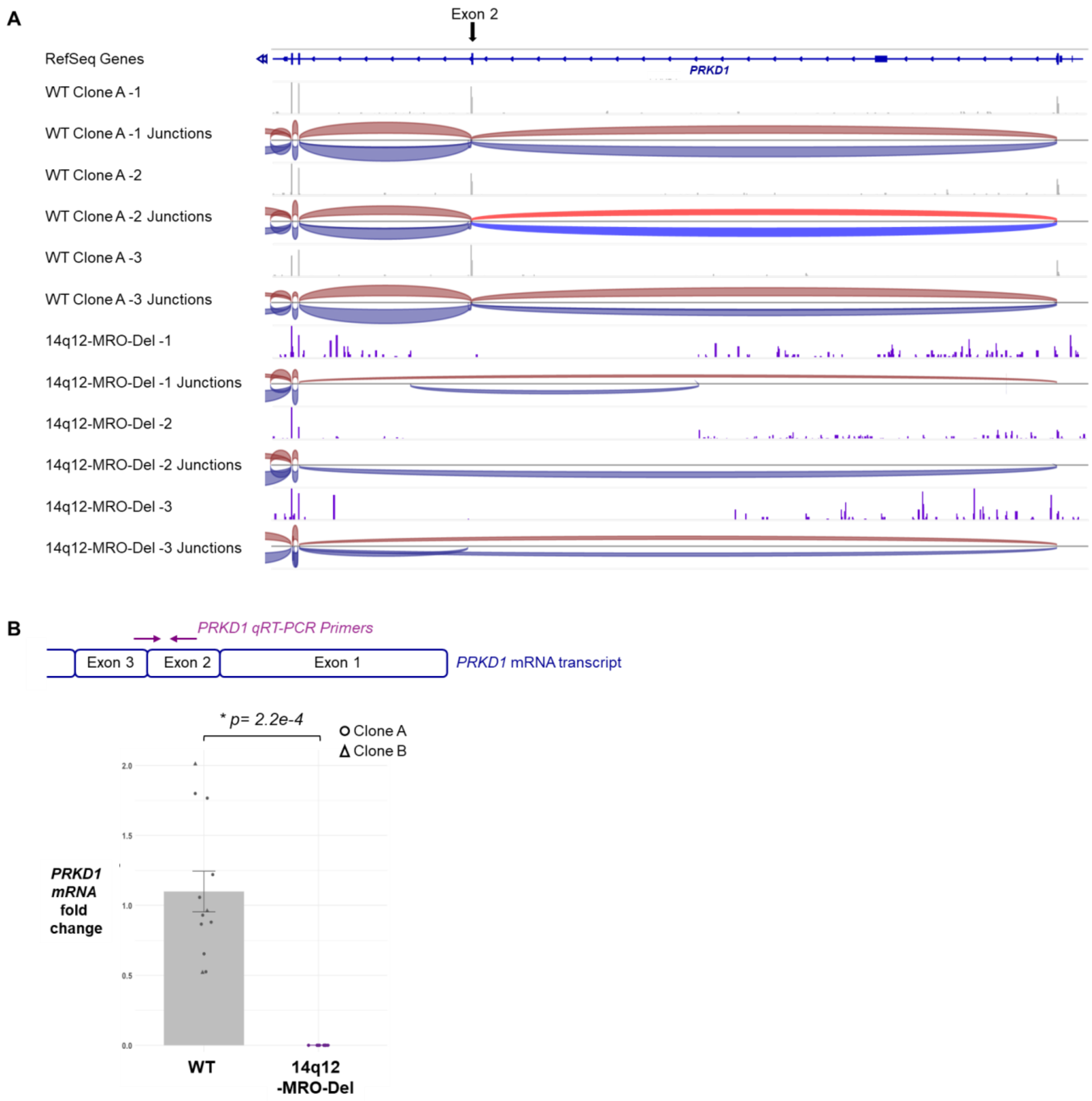


**Supplementary Figure 4.** mRNA analysis of *PRKD1*. (A) IGV plot from RNA-seq of WT and 14q12-MRO-Del HAP1s demonstrating the deletion of exon 2 (arrow) of *PRKD1*, which renders the transcript out of frame, likely subjecting it to nonsense mediated decay. The junction reads (blue and red) confirm skipping of this exon (n=3) (B) *PRKD1* mRNA expression assayed by qRT-PCR in WT and 14q12-MRO-Del HAP1s, with primers spanning *PRKD1* exon 2 as shown. (Replicate count: WT Clone A = 9, WT Clone B = 3, 14q12-MRO-Del = 9). * Two sample T-test was performed for statistical significance.

| **VISTA Enhancer** | **Activity** | **Example embryo (E11.5)** |
| --- | --- | --- |
| Hs1064 | Forebrain, Hindbrain | 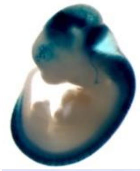 |
| Hs566 | Forebrain | 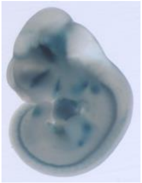 |
| Hs1539 | Hindbrain | 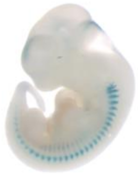 |
| Hs1168 | Facial mesenchyme, Hindbrain, Cranial nerve | 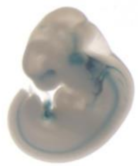 |
| Hs1523 | Branchial arch, Facial mesenchyme, Forebrain, Midbrain, Hindbrain, Eye | 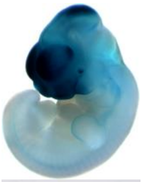 |
| Hs342 | Forebrain | 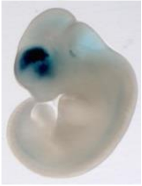 |
| Hs598 | Neural tube | 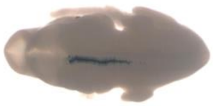 |
| Hs433 | Forebrain, Hindbrain | 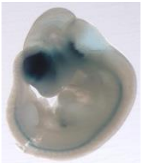 |
| Hs344 | Forebrain | 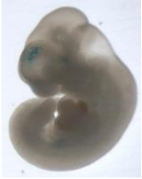 |

**Supplementary Figure 5.** Active VISTA enhancers in the 14q12 regulatory region downstream of *FOXG1* [1].

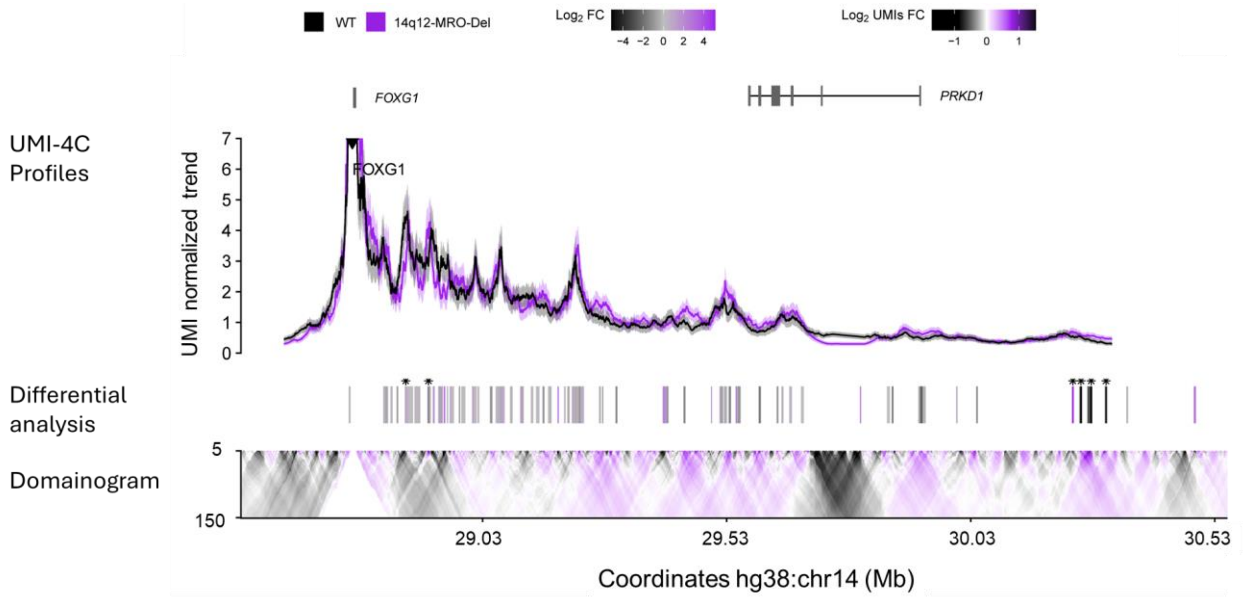


***Genomic intervals of differential UMI-4C contacts between WT and 14q12-MRO-Del**

| Coordinates (hg19) | Coordinates (hg38) | p-value | log2FoldChange |
|---|---|---|---|
| chr14:29,340,992-29,344,417 | chr14:28,871,786-28,875,211 | 0.0021 | 1.13 |
| chr14:29,387,328-29,391,265 | chr14:28,918,122-28,922,059 | 0.0016 | -1.42 |
| chr14:30,707,603-30,710,998 | chr14:30,238,397-30,241,792 | 0.0013 | 5.14 |
| chr14:30,724,104-30,728,011 | chr14:30,254,898-30,258,805 | 0.0015 | -4.98 |
| chr14:30,745,035-30,748,796 | chr14:30,275,829-30,279,590 | 0.0019 | -4.89 |
| chr14:30,775,521-30,779,248 | chr14:30,306,315-30,310,042 | 0.0012 | -5.14 |

**Supplementary Figure 6.** Differential UMI-4C contact analysis. The top panel of UMI-4C profiles represents a smoothed trend of normalized counts of *FOXG1* genomic interactions from WT (black) and 14q12-MRO-Del (purple) samples. The black inverted triangle is directed towards the *FOXG1* viewpoint. The differential analysis panel represents $\log_2$ fold change (FC) of differential UMI-4C contacts in the 14q12 region from WT and 14q12-MRO-Del samples. Asterisks in black indicate significant differences in contacts between samples (FDR adjusted *p*-value $< 0.05$). The bottom panel is a domainogram that illustrates the mean contact intensity $\log_2$ fold changes between WT and 14q12-MRO-Del. * refer to genomic coordinates of the differential UMI contacts where negative log2FC is indicative of increased contacts in the 14q12-MRO-Del line, positive values are loci that have decreased contacts in the 14q12-MRO-Del line (higher in WT).

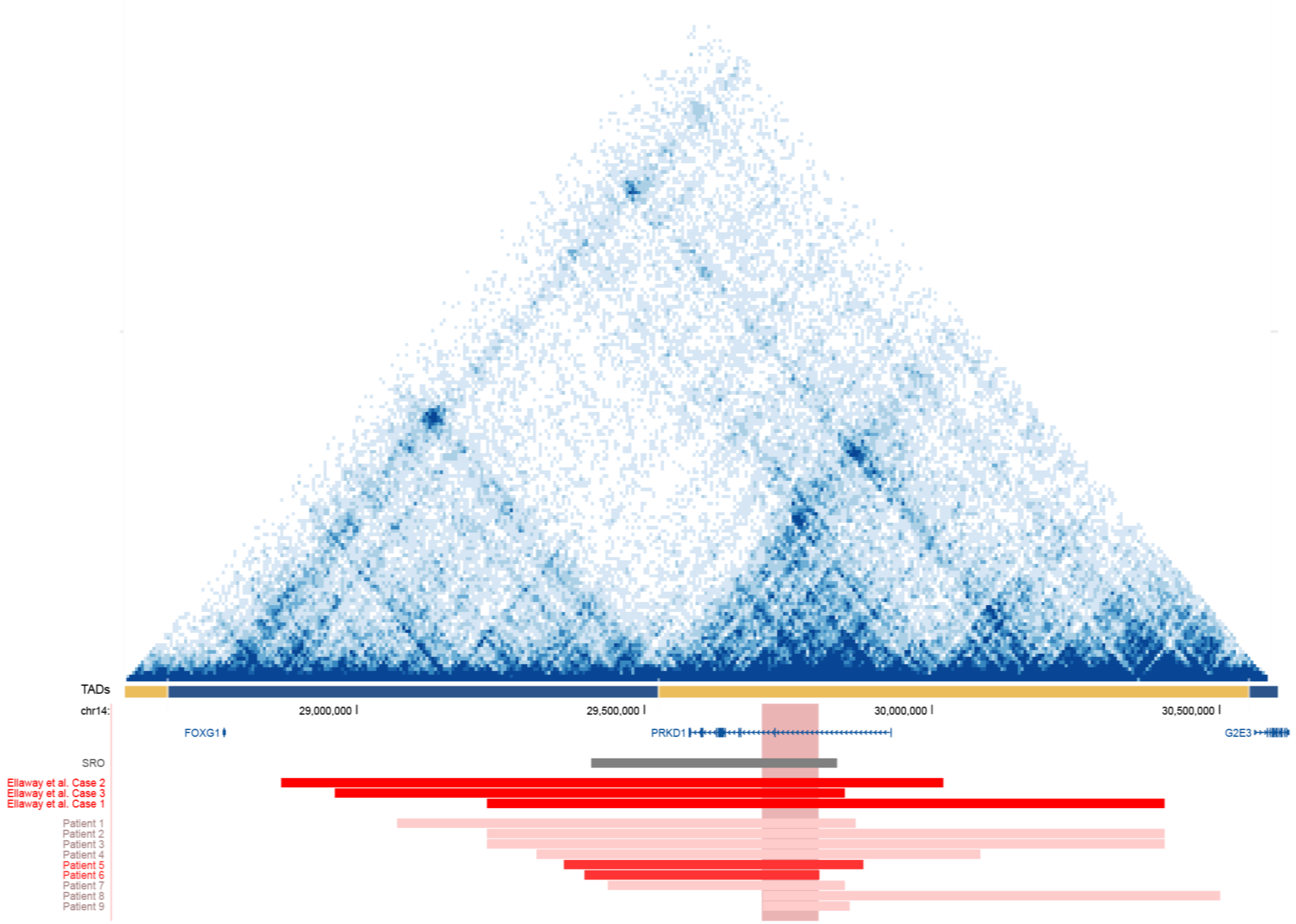


**Supplementary Figure 7.** Deletions downstream of *FOXG1* in the context of native 14q12 chromatin architecture. HAP1 Hi-C data matrix (previously generated by [2]) with TAD annotations reveal the SRO [3] as well as a majority of the 14q12 deletions (this study and [4]), span both the TAD containing *FOXG1* and a neighboring TAD. The MRO (vertical highlight in red) resides in the TAD neighboring *FOXG1*. Hi-C data was visualized using the 3D Genome Browser [5].

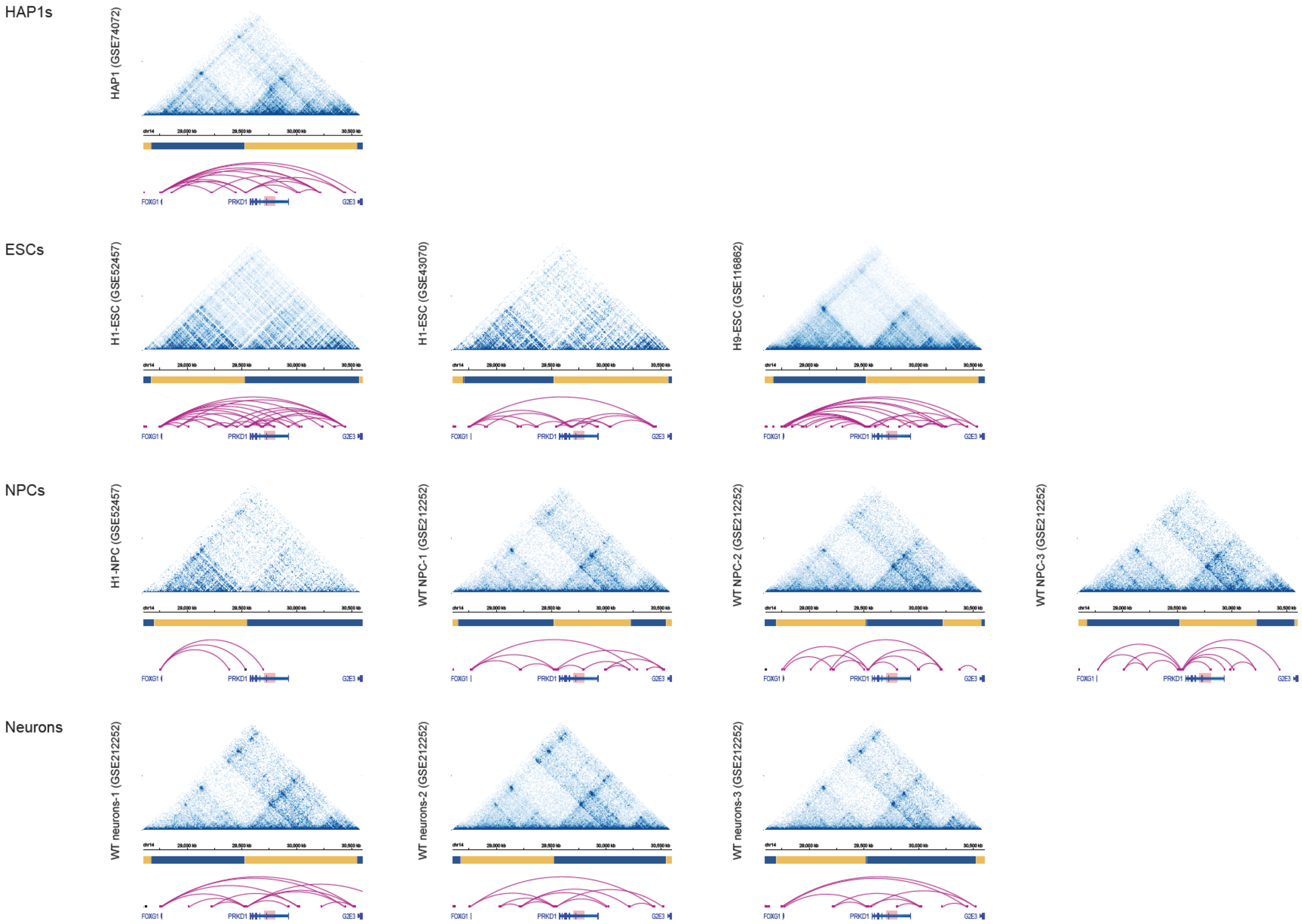


**Supplementary Figure 8.** Native 14q12 chromatin architecture across various cell types. Hi-C data from HAP1s, H1 and H9 (ESCs), neural progenitor cells (NPCs) and NPC-derived neurons reveal a highly conserved *FOXG1* TAD boundary across cell types, with chromatin interactions spanning TAD boundaries. TADs are annotated by alternating horizontal blue and yellow bars. The MRO is highlighted in red. Hi-C data was visualized using the 3D Genome Browser [5].

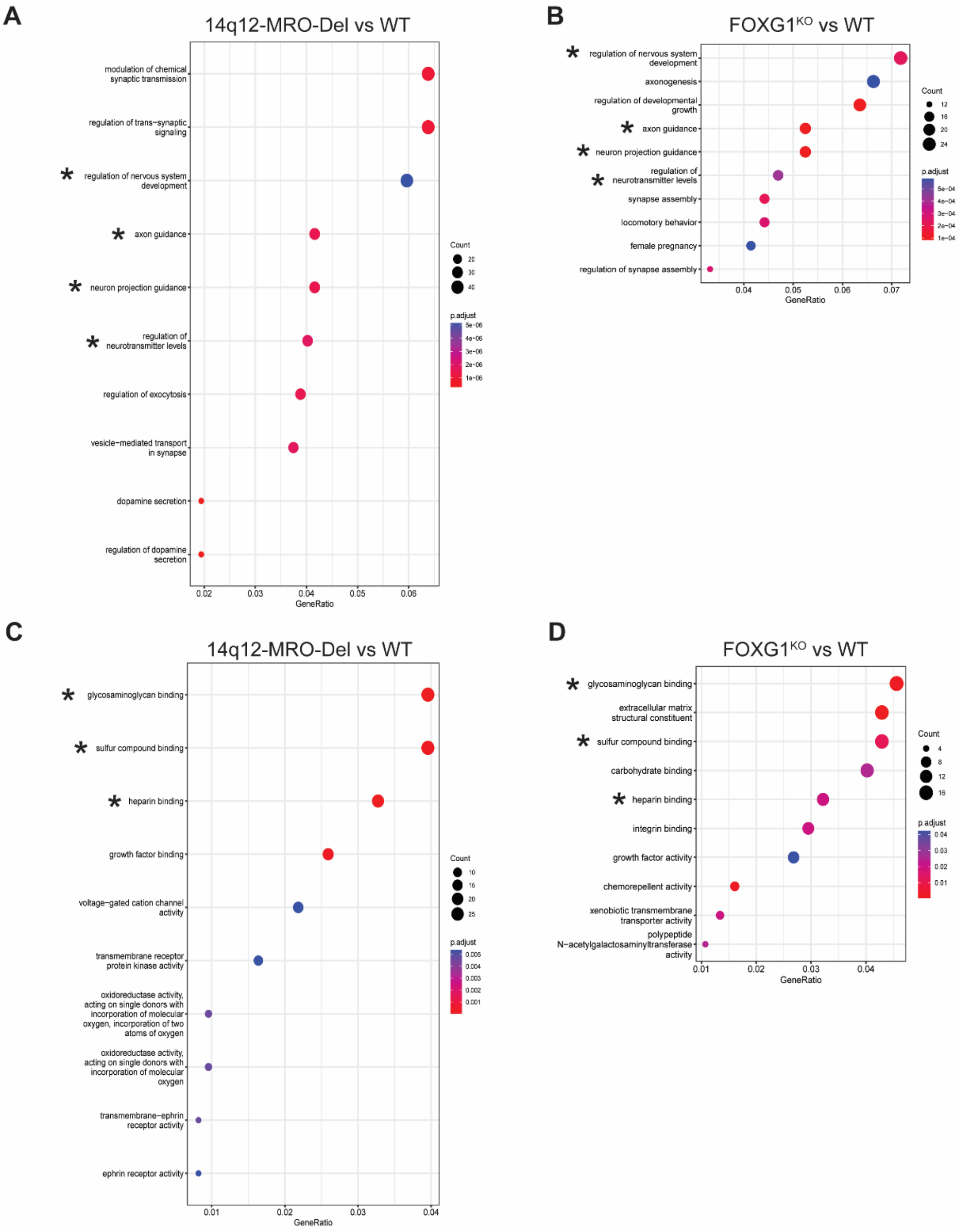


**Supplementary Figure 9.** Functional enrichment (FE) analysis of DEGs. (A-B) Biological pathways and (C-D) molecular functions that are shared between 14q12-MRO-Del vs WT and FOXG1$^{KO}$ vs WT datasets are designated as *

**Supplementary Table 1**. Catalog of gRNAs and oligos

| Protocol | Specific Usage | Primer Sequence (5' to 3') |
|---|---|---|
| CRISPR/ Cas9 gene editing | MRO left breakpoint gRNA | ACATATAGAATGGAAGTTTG |
| | MRO right breakpoint gRNA | CTTCAGCTCACTGCTTACTG |
| | FOXG1$^{KO}$ left breakpoint gRNA | AGCATCACCCAGTCGTCGGG |
| | FOXG1$^{KO}$ right breakpoint gRNA | TCACTTACAGTCTGGTCCCA |
| Genotyping PCRs | MRO breakpoint PCR F | TCATTCTTGCCAAAGTAAAACA |
| | MRO breakpoint PCR R | TGCCCGGCTAATTTTTGTAT |
| | FOXG1$^{KO}$ breakpoint PCR F | GGCCAGACCAGTTACTTTTTCC |
| | FOXG1$^{KO}$ breakpoint PCR R | TTTAGGTTGTTCTCAAGGTCTGC |
| qRT-PCR | *FOXG1* Primer F | CCTGCCCTGTGAGTCTTTAAG |
| | *FOXG1* Primer R | GTTCACTTACAGTCTGGTCCC |
| | *PRKD1* Primer F | CTTTTTCGCCATGACCCTACC |
| | *PRKD1* Primer R | GGAAGCTGACAAGACCACTTCA |
| | *EAR* Primer F | GAGGCTGAGGCAGGAGAATCG |
| | *EAR* Primer R | GTCGCCCAGGCTGGAGTG |
| UMI-4C | *FOXG1* viewpoint US primer | GGGTGGGGAGGATGGAATAA |
| | *FOXG1* viewpoint DS primer | AATGATACGGCGACCACCGAGATCTAC ACTCTTTCCCTACACGACGCTCTTCCG ATCTTGAGTATTCATCAACCGCATTCT |
| | Illumina Universal Primer 2 | CAAGCAGAAGACGGCATACGA |

**Supplementary Table 2.** Catalog of cell lines

| Cell Line | Genotype |
|---|---|
| WT-Clone A (unedited) | *FOXG1*$^{+,}$ MRO$^{+}$ |
| WT-Clone B | *FOXG1*$^{+,}$ MRO$^{+}$ |
| 14q12-MRO-Del | *FOXG1*$^{+,}$ MRO$^{-}$ |
| FOXG1$^{KO}$ | *FOXG1*$^{-,}$ MRO$^{+}$ |

Abbreviations: WT, wildtype; KO, knock-out, Del, deletion

**Supplementary Table 3.** Catalog of antibodies

| Antibody | Experiment | Binding Condition | Product Information |
|---|---|---|---|
| FOXG1 (epitope within the last 100 amino acids) | Western (primary) | 1:1000 overnight at 4°C | Abcam 196868 |
| TAF5 | Western (primary) | 1:1000 overnight at 4°C | Bethyl Labs A303-686A |
| Goat anti-Rabbit IgG H&L (HRP) | Western (secondary) | 1:20,000 1 hr at RT | Abcam 205718 |

**Supplementary Table 4.** Genomic intervals of interest in UMI-4C analysis

| Interval ID | Coordinates (hg19) | Coordinates (hg38) | Size |
|---|---|---|---|
| *FOXG1* promoter bait | chr14: 29,234,251-29,234,254 | chr14: 28,765,045-28,765,048 | 4 bp |
| 14q12-MRO | chr14: 30,173,942-30,273,103 | chr14: 29,704,736-29,803,897 | 99 kb |
| 14q12-MRO-Del | chr14: 30,173,921-30,273,106 | chr14: 29,704,715-29,803,900 | 99 kb |
| SRO | chr14: 29,875,672-30,303,083 | chr14:29406466-29833877 | 427kb |

**Supplementary Table 5.** Genomic intervals of cCREs tested in luciferase assay

| CRE ID | Coordinates (hg38) |
|---|---|
| cCRE1 | chr14:29,736,248-29,737,089 |
| cCRE2 | chr14:29,799,340-29,799,959 |

**Supplementary Data 1.** Differentially expressed genes (DEGs) and direct FOXG1 targets from RNA-seq

**List of items in Source Data file:**

- Fig. 2A. *FOXG1* mRNA Fold Change Values, normalized to WT.
- Fig. 2B. FOXG1 Protein Fold Change Values, normalized to WT. Densitometry data and Western Blots
- Fig. 3. Genomic coordinates of interest including GWAS and MPRA hits.
- Fig. 4B. Luciferase Reporter Assay Fold Change Values, normalized to Empty.
- Supplementary Fig. 1. Normalized transcript per million (nTPM) values for FOXG1 across various cell/tissue types.
- Supplementary Fig. 4B. *PRKD1* mRNA Fold Change Values, normalized to WT.
- Supplementary Fig. 6. Differential genomic interactions between samples from UMICats.

**SUPPLEMENTARY REFERENCES:**


1. Kosicki, M. *et al.* VISTA Enhancer browser: an updated database of tissue-specific developmental enhancers. *Nucleic Acids Res* **53**, (2025).

2. Sanborn, A. L. *et al.* Chromatin extrusion explains key features of loop and domain formation in wild-type and engineered genomes. *Proc Natl Acad Sci U S A* **112**, E6456–E6465 (2015).

3. Allou, L. *et al.* 14q12 and severe Rett-like phenotypes: new clinical insights and physical mapping of FOXG1-regulatory elements. *Eur J Hum Genet* **20**, 1216–1223 (2012).

4. Ellaway, C. J. *et al.* 14q12 microdeletions excluding FOXG1 give rise to a congenital variant Rett syndrome-like phenotype. *European Journal of Human Genetics 2013 21:5* **21**, 522–527 (2012).

5. Wang, Y. *et al.* The 3D Genome Browser: A web-based browser for visualizing 3D genome organization and long-range chromatin interactions. *Genome Biol* **19**, 1–12 (2018).
